# Regulation of extracellular matrix assembly and structure by hybrid M1/M2 macrophages

**DOI:** 10.1101/2020.08.21.261933

**Authors:** Claire E. Witherel, Kimheak Sao, Becky K. Brisson, Biao Han, Susan W. Volk, Ryan J. Petrie, Lin Han, Kara L. Spiller

**Affiliations:** School of Biomedical Engineering, Science, and Health Systems, Drexel University, Philadelphia, PA; Department of Biology, College of Arts and Sciences, Drexel University, Philadelphia, PA; Department of Clinical Sciences and Advanced Medicine, University of Pennsylvania School of Veterinary Medicine, Philadelphia, PA

**Keywords:** macrophage phenotype, hydrogel, fibrous capsule, ECM assembly, fibroblast

## Abstract

Aberrant extracellular matrix (ECM) assembly surrounding implanted biomaterials is the hallmark of the foreign body response, in which implants become encapsulated in thick fibrous tissue that prevents their proper function. While macrophages are known regulators of fibroblast behavior, how their phenotype influences ECM assembly and the progression of the foreign body response is poorly understood. In this study, we used *in vitro* models with physiologically relevant macrophage phenotypes, as well as controlled release of macrophage-modulating cytokines from gelatin hydrogels implanted subcutaneously *in vivo* to investigate the role of macrophages in ECM assembly. Primary human macrophages were polarized to four distinct phenotypes, which have each been associated with fibrosis, including pro-inflammatory M1, pro-healing M2, and a hybrid M1/M2, generated by exposing macrophages to M1- and M2-promoting stimuli simultaneously. Additionally, macrophages were first polarized to M1 and then to M2 (M1→M2) to generate a phenotype typically observed during normal wound healing. Human dermal fibroblasts that were cultured in macrophage-conditioned media upregulated numerous genes involved in regulation of ECM assembly, especially in M2-conditioned media. Hybrid M1/M2 macrophage-conditioned media caused fibroblasts to produce a matrix with thicker and less aligned fibers, while M2 macrophage-conditioned media caused the formation of a more aligned matrix with thinner fibers. Gelatin methacrylate hydrogels containing interleukin-4 (IL4) and IL13-loaded poly(lactic-co-glycolic acid) (PLGA) microparticles were designed to promote the M2 phenotype in a murine subcutaneous *in vivo* model. NanoString multiplex gene expression analysis of hydrogel explants showed that hydrogels with and without drug caused markers of both M1 and M2 phenotypes to be highly expressed, but the release of IL4+IL13 promoted upregulation of M2 markers and genes associated with regulation of ECM assembly, such as *Col5a1* and *Col6a1*. Biochemical analysis and second harmonic generation microscopy showed that the release of IL4+IL13 increased total sulfated glycosaminoglycan content and decreased fibril alignment, which is typically associated with less fibrotic tissue. Together, these results show that hybrid M1/M2 macrophages regulate ECM assembly, and that shifting the balance towards M2 may promote architectural and compositional changes in ECM with enhanced potential for downstream remodeling.

## 1. Introduction

Extracellular matrix (ECM) assembly is a critical and tightly regulated process in tissue repair. Dysfunctional ECM assembly leads to fibrosis, a major cause of morbidity and mortality, which affects tissues as diverse as the liver, kidney, lungs, heart, and skin. The foreign body response, a form of biomaterial-mediated fibrosis in which biomaterials become encapsulated within fibrous tissue and isolated from the rest of the body, impedes the function of many biomaterial implants. The foreign body response is similar to fibrosis observed in other pathologic conditions, such as end-stage kidney and liver disease and chronic autoimmune diseases like scleroderma and rheumatoid arthritis, with hallmarks including excessive collagen deposition, increased matrix stiffness, and highly aligned ECM [1–4]. Strategies to modulate the fibrotic response to biomaterials are in critical need.

The normal response to injury, including the implantation of a biomaterial, involves the recruitment of immune cells, especially macrophages, as well as the proliferation of fibroblasts and their secretion of extracellular matrix (ECM). Macrophages are essential to the wound healing process and possess the ability to modulate fibroblast activities and fate to govern the quality of matrix repair. Key to these diverse functions of macrophages is their phenotype plasticity as wound healing progresses. In the early stages of the response to injury, macrophages primarily assume a pro-inflammatory phenotype, also known as M1; in later stages of healing, macrophages transition into an alternative (M2) phenotype that is associated with resolution of the healing process [5]. However, the extent of the diversity of the M2 population is not known, and numerous M2 subtypes have been described [6]. While the field continues to evolve, the current state of knowledge is that M1 macrophages initiate healing while M2 macrophages promote healing resolution (for review, see [7, 8]). In addition, recent studies employing single cell transcriptomic profiling have shown that macrophages often exist as hybrid phenotypes *in vivo*, simultaneously expressing markers of both archetypal M1 and M2 phenotypes [9]. Because macrophages switch from M1 to M2 during the course of normal wound healing, the generation of a hybrid M1/M2 phenotype in between these phases is likely. However, the roles of hybrid M1/M2 phenotypes in wound healing are not clear.

Fibrosis can occur when wound healing fails to resolve and is generally considered to be pathologically Th2-driven and mediated by M2-type macrophages [10], but both M1 and M2-type macrophages have been associated with fibrosis in clinical studies and animal models. For example, pro-inflammatory cytokines like tumor necrosis factor-α (TNFα), interleukin-1β (IL1β), and IL6, which are secreted at high levels by M1 macrophages, are strongly associated with fibrosis [11–13]. On the other hand, the potent M2-promoting cytokines IL4 and IL13 as well as M2-secreted factors like transforming growth factor-β (TGFβ), platelet-derived growth factor-BB (PDGFBB), CCL17, and CCL22 are also hallmarks of fibrosis [14–16]. Other studies that analyzed macrophages participating in fibrous encapsulation of biomaterials have reported co-expression of both M1 and M2 markers [9, 17–19]. Also, while the M2-promoting cytokine IL4 stimulates the fusion of macrophages into multinucleated foreign body giant cells [20], several recent reports have shown that controlled release of IL4 from biomaterials promoted a significant reduction in fibrous capsule formation around implanted biomaterials [21, 22]. These conflicting reports can be potentially resolved by the increasing understanding that macrophages exist *in vivo* on a spectrum of phenotypes that can rarely be defined as precisely as *in vitro*-polarized M1 and M2 macrophages. However, despite the growing appreciation of the existence *in vivo* of hybrid phenotypes, the role of hybrid macrophage phenotypes in biomaterial-mediated fibrosis is still unknown.

To address these gaps in knowledge, this study took a two-pronged approach to investigate the role of macrophage phenotype in ECM assembly and fibrosis *in vitro* and *in vivo*. First, we examined the effects of physiologically relevant macrophage phenotypes, including hybrid M1/M2 phenotypes, on fibroblast behavior *in vitro* with respect to gene expression, ECM deposition and structure, and matrix stiffness. Macrophages with a hybrid M1/M2 phenotype were prepared by culturing in M1- and M2-promoting stimuli simultaneously. Another unique, hybrid phenotype was prepared by culturing macrophages first in M1-promoting stimuli and then switching to M2-promoting stimuli (M1→M2), a transition that occurs in normal wound healing and which results in an M2-like phenotype with some M1 characteristics [23]. Second, we designed M2-promoting gelatin hydrogels via the controlled release of IL4+IL13 to investigate the effects of shifting the macrophage population towards M2 on ECM assembly within the fibrous capsule in a murine subcutaneous model.

## 2. Methods

### 2.1 Macrophage differentiation and polarization

Primary human peripheral blood monocytes (n=4 donors) were purchased from the University of Pennsylvania Human Immunology Core and differentiated into macrophages through the addition of macrophage colony stimulating factor (MCSF), as previously described [17]. M1 macrophages were prepared via 72 hours of incubation with 20ng/mL MCSF plus 100ng/mL interferon-γ (IFNγ) and 100 ng/ml lipopolysaccharide (LPS); M2 macrophages were prepared with 20ng/mL MCSF, 40 ng/mL IL4, and 20 ng/mL IL13; hybrid M1/M2 macrophages were prepared with the same concentrations of IFNγ, LPS, IL4, and IL13, given simultaneously; and finally M1→M2 macrophages were prepared by first culturing macrophages in M1-promoting stimuli (IFNγ and LPS) for 24 hours, washing cells with phosphate buffered saline (PBS), followed by M2-promoting stimuli (IL4 and IL13) for an additional 48 hours (**Fig. 1A**). Conditioned media without the presence of polarizing cytokines was generated at the end of the polarization period by washing the cells once in PBS and then incubating in basal media for 24 hours prior to storage at −80°C until later use.

**Figure 1.**
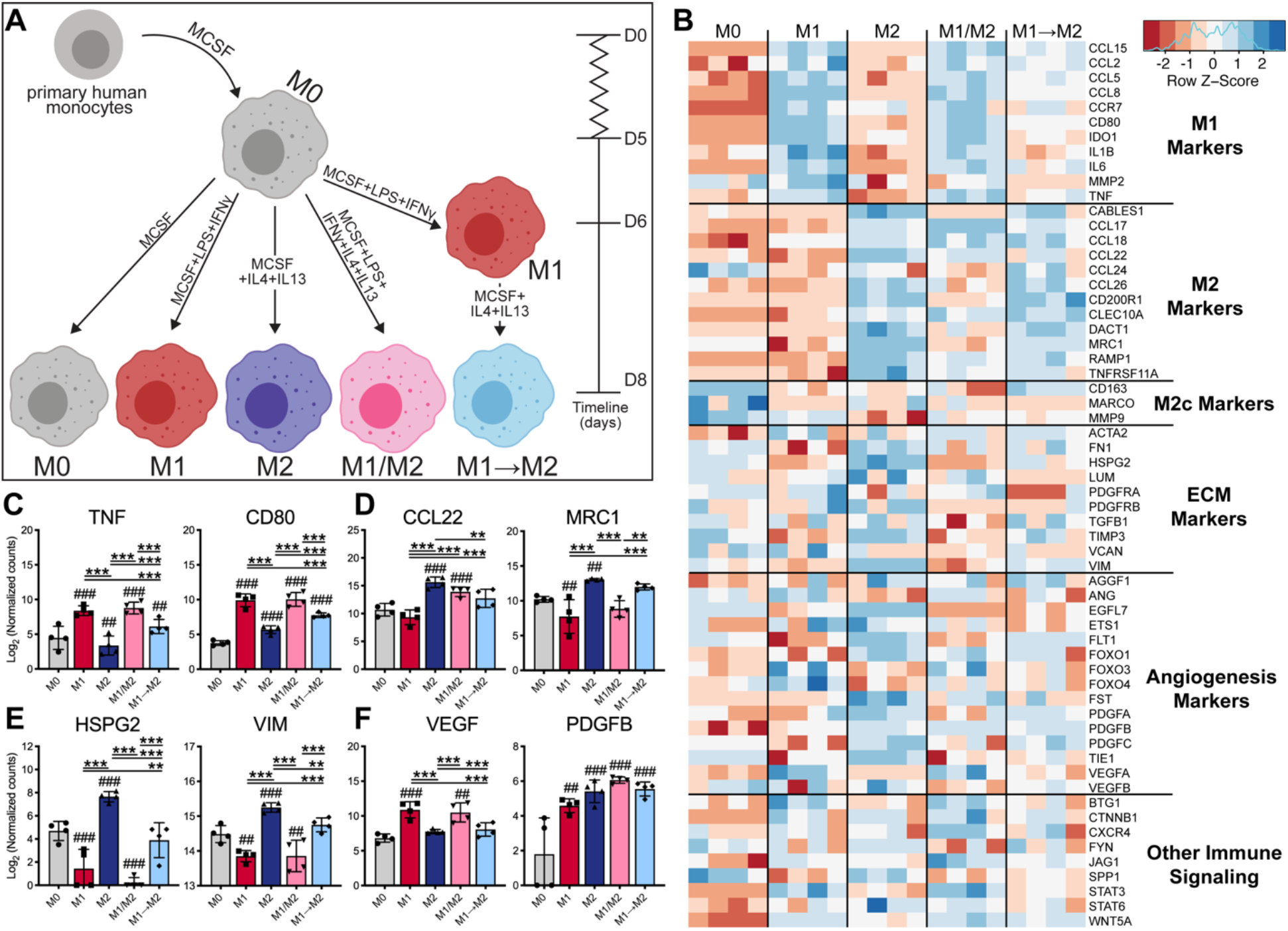
M2 macrophages upregulated genes associated with ECM assembly in vitro. A) Schematic of primary human monocyte to macrophage culture and subsequent polarization *in vitro*. B) Heatmap of all expressed genes represented as the row Z-score of log-transformed normalized data (normalized to negative and positive controls and housekeeping genes, GAPDH and TBP). Genes that were not expressed above background levels were: BGN, COL1A1, COL3A1, COL5A1, CCN2, CXCL12, DCN, FGF2, IGF1, VEGFC. Two representative differentially expressed genes from the categories related to C) M1, D) M2 markers, E) ECM, and F) Angiogenesis markers. All genes are plotted in **Suppl. Figs. 1-6**. Data represented as mean ± standard deviation (SD), n=4 donors. **, ## p<0.01, ***, ### p<0.001, where # symbols indicate statistical significance compared to M0 control.

### 2.2 Fibroblast culture and matrix analysis

Primary human dermal fibroblasts (American Tissue Culture Collection) were maintained in basal media: Dubecco’s Modified Eagle Medium (DMEM), supplemented with 10% fetal bovine serum and 1% penicillin-streptomycin-glutamine, before transferring to glutaraldehyde-crosslinked gelatin coated glass-bottom 35mm petri dishes. After cells were >80% confluent, macrophage conditioned media was added at a 1:1 ratio of basal fibroblast media to macrophage conditioned media. Media was changed every other day for 14 days. Fibroblasts together with any deposited ECM were either immediately processed for gene expression analysis by lysing the cells in 100μL Buffer RLT (Qiagen, RNeasy Mini kit), or matrices were denuded of cells by treating with 20mM ammonium hydroxide + 0.5% TritonX-100 in PBS for a few seconds, followed by two washes with PBS. Matrices were then cleared of any residual DNA by treating with DNAse at 10 U/mL at 37°C for 30 minutes, followed by two washes of PBS. Matrices were fluorescently labeled using Alexa Fluro™ 647 NHS Ester (Thermo Fisher) reconstituted in dimethyl sulfoxide (DMSO) and diluted to 10μg/ml in PBS supplemented with penicillin-streptomycin to a concentration of 10mg/mL. Confocal microscopy of resultant matrices was conducted on a Zeiss LSM 700 using the 63x/1.4 oil objective at either 0.5x digital zoom or 2.0x digital zoom (for image analysis via ImageJ) to visualize matrix and to determine matrix thickness (measured by recording the z-stack height). High magnification images were taken of each matrix, with n=3 images per group per biological replicate (n=3 human donors of macrophages). Each image was imported into ImageJ and compressed into a 2D maximum projection of maximum intensity. The images were then processed for DiameterJ Segmentation, and SuperPixel data was subsequently used and normalized against unit/pixel to obtain fiber diameters [24]. To measure alignment of the fibers in the matrices, the von Mises concentrations, κ were calculated as we previously described [25]. Briefly, for every image, DiameterJ reports every detectable fiber angle between the primary axis and a line parallel to the x-axis of the image. All reported angles from each technical image replicate were pooled and analyzed, where κ indicates how closely the individual fibril orientation angles cluster around the average angle; higher κ suggests a higher degree of fiber alignment.

### 2.3. Gene expression analysis via qRT-PCR or NanoString

Total RNA was extracted from cells using a Qiagen RNeasy kit according to the manufacturer’s instructions. For quantitative RT-PCR (qRT-PCR), RNA was reverse-transcribed after DNAse treatment and processed using SYBR green and primers synthesized by Life Technologies (**Suppl. Table 1**) as we have previously described [26]. Gene expression of the targeted genes was determined by normalizing genes to a reference gene, *b-actin* (2^-ΔCt^ method). NanoString multiplex gene expression analysis was performed using 100ng RNA and custom-designed panels of 72 genes corresponding to human (for *in vitro* studies) or murine (for *in vivo* studies) genes related to macrophage phenotype, fibrosis, and angiogenesis (**Suppl. Tables 2 and 3**). NanoString data were first normalized to internal positive and negative controls and subsequently normalized to the geometric mean of two housekeeping genes GUSB and TBP, as recommended by the manufacturer. Heatmaps were generated in R with *heatmap.2* function in *gplots* package using the row Z-score of log-transformed normalized data.

### 2.4. Nanoindentation via atomic force microscopy

Atomic force microscopy (AFM)-based nanoindentation was carried out on the fibroblast-derived matrices grown on glass coverslips in 1X PBS at room temperature using a Dimension ICON AFM (Bruker Nano, Santa Barbara, CA) and a colloidal spherical tip (radius R ≈ 5μm). The spherical tip was prepared by manually gluing a polystyrene microsphere (PolySciences, Inc. Warrington, PA) onto a tipless silicon nitride cantilever (cantilever C, nominal spring constant k ≈ 0.6N/m, HQ:NSC36/tipless/Cr-Au, NanoAndMore USA, Watsonville, CA) with M-Bond 610 epoxy (SPI Supplies, West Chester, PA) using the same Dimension ICON AFM. At each indentation location, the probe tip was programmed to indent the sample at a 10μm/s constant z-piezo displacement rate (approximately equals the indentation depth rate) up to a ~10nN maximum indentation force (corresponding to 1 to 2μm maximum indentation depths for each sample). For each specimen, indentation was performed on relatively flat regions (surface roughness <40nm for 5μm × 5μm contact mode imaging) to minimize the impact of surface roughness. At least 10 different indentation locations were tested on each sample. For each indentation curve, the cantilever deflection (in volts) and z-piezo displacement (in μm) were converted to an indentation force (in nN) and depth (in μm) through calibrating the cantilever deflection sensitivity (nm/V) by indenting on a hard mica substrate and a spring constant (nN/nm) via thermal vibration [27]. The initial tip-sample contact point was determined via an algorithm reported previously for soft materials in the absence of attractive interactions [28]. The loading portion of the curve at each location was fit to the finite-thickness corrected Hertz model via least-squares linear regression to calculate the effective indentation modulus at the given indentation rate [29].

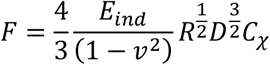

where F and D are the indentation force and depth, respectively, R is the colloidal tip radius, and v is Poisson’s ratio (v = 0.5, similar to other cell-derived matrices) [30]. The correction factor *C_χ_* is 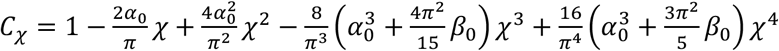 where 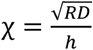, *h* is the thickness of the sample. The constants *α*_0_ and *β*_0_ are functions of the Poisson’s ratio v. In this model, the polystyrene spherical colloid was assumed to have an infinite modulus (~4 GPa) compared to that of the samples, therefore does not deform during the process [31].

### 2.5. M2-promoting hydrogel fabrication

Poly(lactic-co-glycolic acid) (PLGA) microparticles were loaded with IL4 and IL13 and embedded within gelatin methacrylate (GelMA) hydrogels. First, IL4 and IL13 were encapsulated in poly(lactic-co-glycolic acid) (PLGA) microparticles using a double emulsion technique. PLGA (100mg, Sigma Aldrich, Resomer^®^ RG 504 H) was dissolved in 2mL of dichloromethane (DCM) in a glass scintillation vial. Lyophilized murine IL4 and IL13 (Peprotech, Rocky Hill, NJ) were first resuspended in 0.1% fetal bovine serum to a concentration of 1mg/mL, setting aside the appropriate aliquots necessary to fabricate Low (0.5μg IL4 and 0.25μg IL13), Medium (10μg IL4 and 5μg IL13), and High (40μg IL4 and 20μg IL13) doses of IL4an d IL13. Cytokine volumes were pre-mixed and then brought up to a final volume of 200μL with 2% poly(vinyl alcohol) (PVA) (MW 89,000-98,000, 99+% hydrolyzed Sigma Aldrich, St. Louis, MO). These aqueous drug solutions were then added dropwise using a micropipetter to the PLGA-DCM mixture and immediately homogenized at approximately 15,000rpm with a hand-held homogenizer for two minutes. This primary emulsion was then added dropwise to 30mL (in a 50mL glass beaker) of 2% PVA, which was also homogenized at approximately 15000rpm with a hand-held homogenizer for 90 seconds, manually moving the beaker around to ensure thorough mixing. A 0.5 inch stir bar was added to the beaker and the mixture was allowed to stir at approximately 400rpm at room temperature overnight to allow DCM to evaporate out of solution. Particles were subsequently washed with three volumes of 20mL with deionized water in pre-weighed 50mL conical tubes, centrifuging particles at 3500xg for 5 minutes. After the final wash, particles were resuspended in a small volume of water and frozen at −80°C for at least 30 minutes before being lyophilized for at least 24 hours. Conical tubes were subsequently weighed to obtain mass yield of particles, which was typically around 70-90mg of particles for each batch.

To facilitate visualization of PLGA microparticles and proteins within the particles under confocal microscopy, IL4 was pre-labeled with Alexa Fluor™ 405 NHS Ester (Thermo Fisher) and IL13 pre-labeled with Alexa Fluor™ 647 NHS Ester (Thermo Fisher) according to manufacturer’s instructions. Briefly, dyes were reconstituted with anhydrous DMSO to a concentration of 10mg/mL. IL4 and IL13 (Peprotech, Rocky Hill, NJ) were dissolved in 0.1M sodium bicarbonate, pH 8.3. 2-5μL of dye was added to each protein tube depending on concentration and allowed to incubate for 1 hour at room temperature. Proteins were dialyzed using a 10K MWCO membrane (Thermo Scientific™ Slide-A-Lyzer™ MINI device) overnight in 1L of 0.1M sodium bicarbonate, on a stir plate stirring at ~100rpm protected from light. Proteins were used within 24 hours to fabricate PLGA particles. To visualize PLGA particles, 50μL of reconstituted tetramethylrhodamine (TRITC) (at a concentration of 10mg/mL) was also loaded into the aqueous phase of the primary emulsion.

Approximately 30-35mg of dry particles were weighed and resuspended in ~1.5mL of pre-warmed (to 37°C) 0.5% lithium phenyl-2,4,6-trimethylbenzoylphophinate (LAP) (Allevi, Philadelphia, PA) in PBS (representing approximately 2% w/v particle solution). This volume was mixed 1:1 with sterile, pre-warmed (to 37°C) 20% GelMA (Allevi, Philadelphia, PA), resulting in a final concentration of 10% GelMA, 1% particles, and 0.25% LAP. Matrices were cast in 35mm petri dishes and crosslinked for 30 minutes, flipping the dishes over halfway through, via exposure to visible light. Individual hydrogel samples were then biopsy-punched using 7mm biopsy punches.

### 2.6 Hydrogel characterization and release studies

The diameters of PLGA microparticles were analyzed using a Coulter Multisizer 4 and a 100μm aperture tube via 25000 events per sample. Encapsulation of fluorescently labeled IL4 and IL13 was confirmed using confocal microscopy, and distribution of TRITC-labeled microparticles were visualized using confocal microscopy. *In vitro* release studies were conducted to measure the encapsulation and release of IL4 and IL13. Hydrogels were placed in 0.5 mL of basal media (RPMI 1640 + 10% fetal bovine serum + 1% penicillin-streptomycin) and maintained on a heated (37°C) orbital shaker rotating at 100rpm, with release samples collected and replaced with fresh media at 1,2, 4, 7, 14, and 21 days. Hydrogel release samples were analyzed for IL4 and IL13 concentrations using ABTS sandwich ELISAs according to manufacturer’s instructions (Peprotech, Rocky Hill, NJ).

### 2.7 Murine subcutaneous implantation model

All animal experiments followed federal guidelines and were conducted under a protocol approved by Drexel University’s Institutional Animal Care and Use Committee. Hydrogels were immediately prepared prior to implantation to ensure minimal drug loss and matched with *in vitro* benchtop release studies. Subcutaneous biomaterial implantation was performed by a creating a single dorsal incision between the shoulder blades of n=3 12-week-old C57BL/6 male mice (pilot/dose-response study) and n=10 males and n=14 females (follow-up study), where Halsted forceps were used to create pockets to insert biomaterials underneath the skin. In the pilot study, four hydrogels (each containing a different dose of IL4+IL13) were implanted per mouse, while in the follow-up study only two hydrogels of the same kind (Blank or IL4+IL13) were implanted per mouse. Mice were anesthetized for surgery using 1.5-5% isoflurane, received a subcutaneous injection of buprenorphine (0.1mg/kg) prior to being shaved, cleaned with ethanol and iodine before beginning surgery. Animal wounds were secured with a single wound clip and animals were allowed to recover in a dark cage on a heating mat. Mice were housed together and monitored daily for the first three days, and then weekly, for body mass and behavioral changes.

Biomaterials were explanted after 3 and 21 or 22 days *in vivo*, as specified. Samples for gene expression were immediately placed in TRizol reagent, samples for histological analysis were placed in 10% neutral buffered formalin (4% paraformaldehyde), and samples that were to be processed for flow cytometric analysis were placed inside empty tubes before adding 0.5-1mL of 2mg/mL collagenase type 4 once all samples were explanted (approximately 30 minutes); samples incubated in collagenase for 15-30 minutes at room temperature during transport back to the lab, before continuing with the flow cytometry staining protocol. Of note, samples processed for histology were explanted to ensure dermis-biomaterial interface remained intact to ensure accurate measurement of fibrous capsule thickness.

### 2.8 Flow cytometry

Following collagenase treatment, samples were passed through a 70μm cell strainer to filter out any large debris and pelleted in pre-warmed RPMI 1640 media (Gibco). Cell pellets were washed with FACS Buffer (PBS + 0.5% fetal bovine serum + 1% 4-(2-hydroxyethyl)-1-piperazineethanesulfonic acid (HEPES) buffer) before being transferred to a 96-well U-bottom plate. Cells were divided to provide positive staining sample, isotype controls, and an unstained control. Samples were first blocked with FcR block (1:200 dilution in FACS Buffer) for 15 minutes at 4°C, before staining for live dead for 30 minutes at 4°C, followed by primary antibody stain for the following leukocyte markers: CD45 (APC, BioLegend catalog# 103111), CD11b (PE, BioLegend catalog# 101207), CD68 (PE/Cyanine 7, BioLegend catalog# 137015), and F4/80 (APC/Cyanine 7, BioLegend catalog# 123117). All antibodies were diluted 1:200, for 20 minutes at 4°C. Samples were thoroughly washed and fixed prior to analysis using BD LSR II flow cytometer.

### 2.9 Histological analysis

Samples in 10% neutral buffered formalin were incubated at 4°C on a rotator for 24 hours before being washed with PBS three times and stored in 70% ethanol for approximately 2 hours before processing and embedding in paraffin wax. Samples were sectioned at 4-5μm, placed on pre-warmed glass slides and baked at 37°C overnight before staining with hemotoxylin and eosin (H&E), Masson’s Trichrome (Azer Scientific, catalog# ES3398), Alcian Blue (pH 1.0) (Azer Scientific, catalog# S3398) and mounting with Cytoseal-60 (Thermo Scientific).

Optical microscopy was performed on an Olympus BX51 using a 4x objective. Images were imported into ImageJ, stitched together using the pairwise stitching plugin [32], subsequently imported into Adobe Photoshop 2020 for white balancing and cropping. Images were reimported into ImageJ to trace and measure the fibrous capsule area using the Measure tool. Briefly, the fibrous capsule on one side of the biomaterial connected to underside of the dermis was manually traced, reporting the area and perimeter. In order to determine the average width of the capsule, it was assumed that the fibrous capsule was generally rectangular in shape; thus, perimeter, P = 2L + 2w, L = length, w = width, and area, A = Lw. The width w and length L can then be recovered as roots of the quadratic x^2^ - (P/2)x + A; in particular, we estimate w = (P - (P^2^ - 16A)^1/2^)/4.

### 2.10 Biochemical analysis: orthohydroxyproline and dimethylmethylene blue assays

Biochemical content of total collagen and sulfated glycosaminoglycans (sGAGs) were measured using an orthohydroxyproline (OHP) assay [33] and a dimethylmethylene blue assay [34], respectively. Briefly, for both assays, tissues were lyophilized, dry masses were recorded, and samples were digested in a papain solution at 60°C overnight. For the OHP assay, hydrolyzation was performed by adding a small volume of concentrated NaOH solution to each sample and autoclaving samples at 120°C for 10 minutes. To form and measure the chromophore, citric acid was added to each sample and standard and mixed with chloramine-T solution. Samples were incubated at room temperature for 20 minutes followed by the addition of aldehyde/perchloric acid solution, vortexed and incubated at 65°C for 15 minutes. Samples were then added to a 96-well plate in duplicate to measure absorbance at 550nm. For the DMMB assay, following papain digestion, 200μL of DMMB reagent (16mg DMMB, 3.04g glycine, 1.6g NaCl, and 95mL of 0.1M acetic acid, passed through 0.45μm filter) was added to 20μL of sample in a 96-well plate in duplicate, shaken for 5 seconds, and measured absorbance at 525nm. Chondroiton-6-sulfate was used to generate a standard curve.

### 2.11 Second harmonic generation imaging

SHG (backwards) imaging of the fibrillar collagen was acquired using a Leica SP8 confocal/multiphoton microscope (Leica Microsystems, Inc., Mannheim, Germany), as previously described [35, 36]. Tile scanned images were taken that contained the entire length of the fibrous capsule under the dermis (10 z-stacks, 1 μm each). For analysis, max projection regions of interest containing the fibrous capsule but not the overlaying dermis or other tissue (fat and skeletal muscle) were identified. To analyze the collagen fiber qualities (number, straightness, width and length) CT-FIRE was used on images with autofluorescence subtracted from the original SHG signal, as previously [35, 36]. The CT-FIRE program identified all fibrillar collagen fibers within the regions of interest, and analyzed each fiber for length, width, and % of fibers that were straight [35–37]. For alignment, a fast Fourier transform powerplot of the fibrillar collagen signal was analyzed [38].

### 2.12 Statistical Analysis

All statistical analysis was performed in GraphPad Prism v8.4. *In vitro* NanoString gene expression analysis and *in vivo* qRT-PCR (pilot/dose dependent study) data were analyzed using a mixed effects model followed by a multiple comparisons test controlling the false discovery rate (0.01 for in vitro data sets, 0.05 for in vivo data set) using a two-stage linear step up correction method of Benjamini, Krieger, and Yekutieli [39, 40]. NanoString gene expression analysis data of *in vivo* (repeated, separate animals study) were analyzed using an ordinary two-way ANOVA followed by a multiple comparisons test controlling the false discovery rate using a two-stage linear step up correction method of Benjamini, Krieger, and Yekutieli. Biochemical, histological, and second harmonic generation data were analyzed using unpaired, two-tailed, student’s t-tests with a Welch’s correction. Principal component analysis was conducted using R Studio using the *prcomp* function and plotted in GraphPad Prism v.8.4. All data are represented as mean ± standard deviation (SD) and sample sizes are noted in each figure legend.

## 3. Results

### 3.1. Preparation of hybrid macrophage phenotypes

Primary human macrophages were polarized *in vitro* to M1 or M2 phenotypes, and two unique hybrid phenotypes: M1/M2, which was generated through the simultaneous addition of both M1- and M2-promoting stimuli, and M1→M2, which was generated by switching macrophages from M1-to M2-promoting stimuli, and compared to an unactivated (M0) control (**Fig. 1A**). Multiplex gene expression analysis confirmed that M1 macrophages expressed the highest levels of typical M1 markers and M2 macrophages expressed the highest levels of typical M2 markers (**Fig. 1B-D**). M1/M2 macrophages expressed both M1 and M2 markers, although with lower levels of M2 markers compared to the M2 group. M1→M2 macrophages expressed high levels of M2 markers and low levels of M1 markers, although some M1 markers remained highly expressed. All four phenotypes expressed varying levels of genes associated with ECM assembly, angiogenesis, and other immune signaling pathways. Of note, M2 macrophages expressed the highest levels of HSPG2, which encodes perlecan, a heparan sulfate proteoglycan, and VIM, which encodes vimentin, a cytoskeletal filament protein that is typically considered a fibroblast marker (**Fig. 1E**). The hybrid M1/M2 group expressed the highest levels of VEGFA, which encodes the highly angiogenic protein vascular endothelial growth factor, as well as the highest levels of PDGFB, which encodes platelet-derived growth factor-BB, a major growth factor involved at later stages of angiogenesis and which also plays a role in fibrosis (**Fig. 1F**) [15]. All additional genes plotted individually are shown in **Suppl. Figs. 1-6.**

### 3.2 Effects of macrophage-conditioned media on fibroblast behavior

Conditioned media from the different macrophage phenotypes had significant effects on fibroblast gene expression (**Fig. 2A, B**). In particular, M2 macrophage-conditioned media caused fibroblasts to express the highest levels of the genes encoding the matrix proteins collagen III (COL3A1), collagen V (COL5A1), fibronectin (FN1), perlecan (HSPG2), and versican (VCAN), which are each involved in the regulation of ECM assembly (**Fig. 2A**). M2 macrophages also induced fibroblasts to express the highest levels of genes involved in angiogenesis, including the genes encoding PDGFA, VEGFA, DACT1, and the tyrosine kinase protein FYN (**Fig. 2B**). Further functional analysis was performed on fibroblast-derived matrices, which were denuded of cells and fluorescently labeled to aid in the visualization via confocal microscopy (**Fig. 2C, Suppl. Fig. 7**). Macrophage-conditioned media did not cause significant changes in overall matrix thickness (**Fig. 2D**), but M1- and M2-conditioned media caused reductions in average fiber diameter compared to the control and M0 groups, while M1/M2-conditioned media caused an increase in fiber diameter compared to the M0 group (**Fig. 2E**). Conditioned media from M2 and M1→M2 macrophages caused increases in von Mises concentration (κ) compared to the control, indicating significantly more aligned matrix fibrils, while the hybrid M1/M2 group caused a significant decrease in κ compared to the control and M0 groups, indicating that matrix fibrils were less aligned (**Fig. 2F**). Finally, significant differences in matrix stiffness were not noted (**Fig. 2G**).

**Figure 2.**
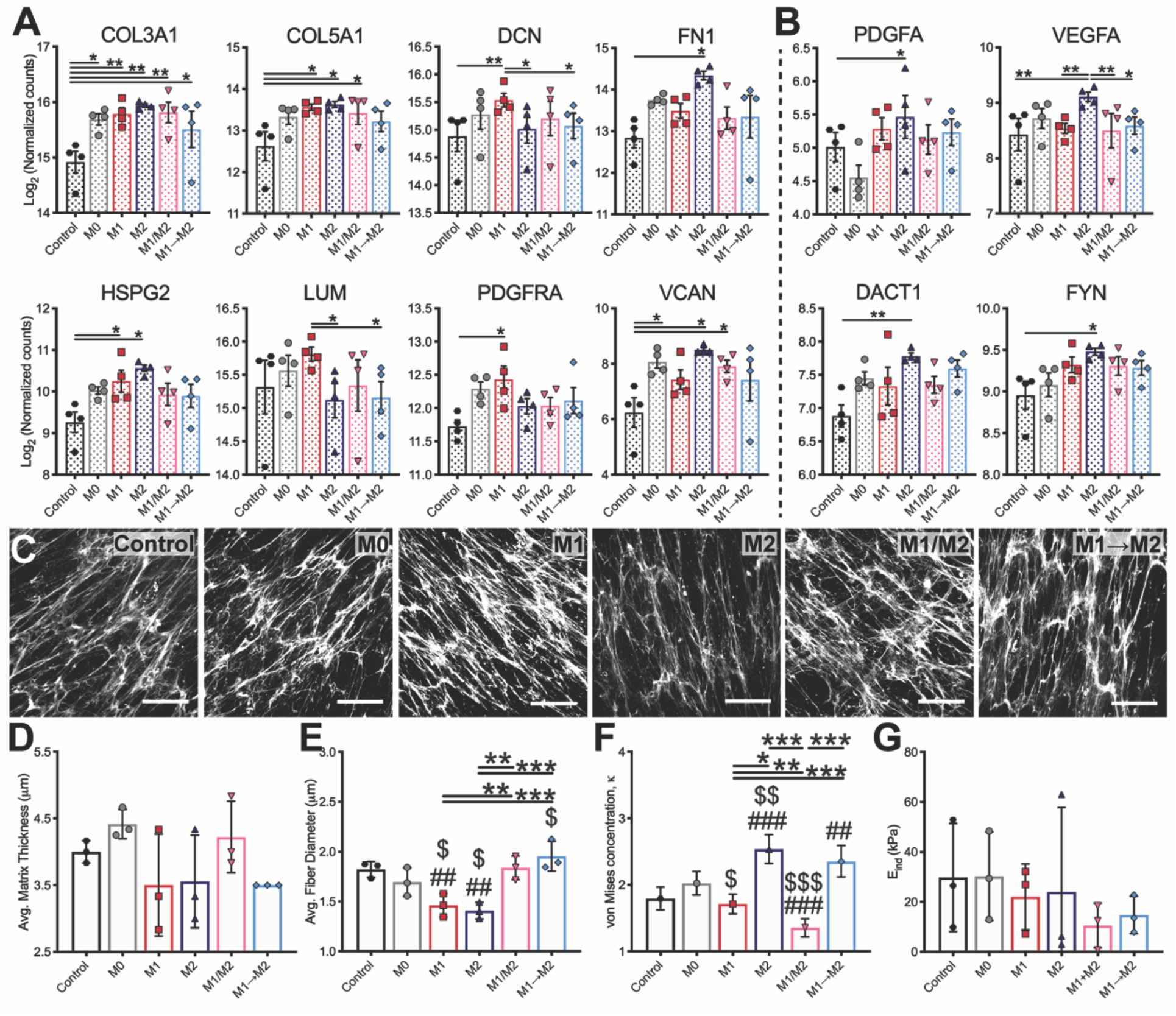
Macrophage-conditioned media affects gene expression and ECM assembly by fibroblasts in vitro. A) Differentially expressed ECM-related and B) angiogenesis-related genes of fibroblasts cultured with primary macrophage-derived conditioned media for 14 days analyzed via NanoString, n=4 primary monocyte donors. C) Images of fibroblast-derived matrix after 14 days via maximum projections of confocal microscopy images. Image analysis (n=3 technical replicates per donor, n=3 donors) was used to determine D) matrix thickness, E) average matrix fiber diameter, and F) von Mises concentration analyzing matrix alignment. G) Indentation modulus of decellularized fibroblast-derived matrices, measured via atomic force microscopy nanoindentation. Each data point represents the average of n=10 experimental indents for each matrix per condition for each biological replicate (n=3 donors). Data represented as mean ± SD, with the exception of von Mises concentration, which is shown as κ with upper and lower limits of a 95% confidence interval. *, $ p<0.05, **, ## p<0.01, ***, ### p<0.001, where # indicates significance compared to the media-only fibroblast control, and $ indicates significance relative to fibroblasts cultured with M0 macrophage-conditioned media.

### 3.3 Effects of IL4+IL13 release on gene expression in vivo

To examine how shifting macrophage phenotype towards M2 affected ECM assembly *in vivo*, gelatin-methacrylate hydrogels were prepared with homogenous distribution of IL4+IL13-loaded PLGA microparticles of approximately 7μm in diameter (**Fig. 3A-E**). Release profiles of IL4 and IL13 were characterized by high burst releases, followed by sustained release for up to 21 days *in vitro* (**Fig. 3F**). The high dose microparticles released approximately 150 ng of IL4 and 100ng of IL13 per hydrogel, while the medium dose released 25 ng IL4 and 8ng IL13, and the low dose released 0.6ng IL4 and 0.3ng IL13 (complete release data shown in **Suppl. Table 4)**. Flow cytometric analysis of the cell population surrounding the hydrogels after 3 or 21 days of subcutaneous implantation in mice showed that approximately 50-60% of the cells were myeloid-derived at day 3, while 75-80% were myeloid-derived at day 21 (**Suppl. Fig. 8**). At 21 days, gene expression analysis of the surrounding cells showed dose-dependent increases in several genes involved in ECM assembly, including the genes that encode connective tissue growth factor (*Ccn2*), decorin (*Dcn*), biglycan (*Bgn*), collagen I (*Col1a1*), collagen III (*Col3a1*), and collagen V (*Col5a1*), although expression of these genes was generally only statistically significant for the high or medium doses of IL4+IL13 compared to the blank control group (**Fig. 3H**). Expression of M1 (**Fig. 3I**) and M2 (**Fig. 3J**) markers were generally unaffected by IL4+IL13 release at this time point, except for significant increases in the M1 marker *Cd38* at the medium and high doses. Additional genes that were not differentially expressed are shown in **Suppl. Fig. 9**.

**Figure 3.**
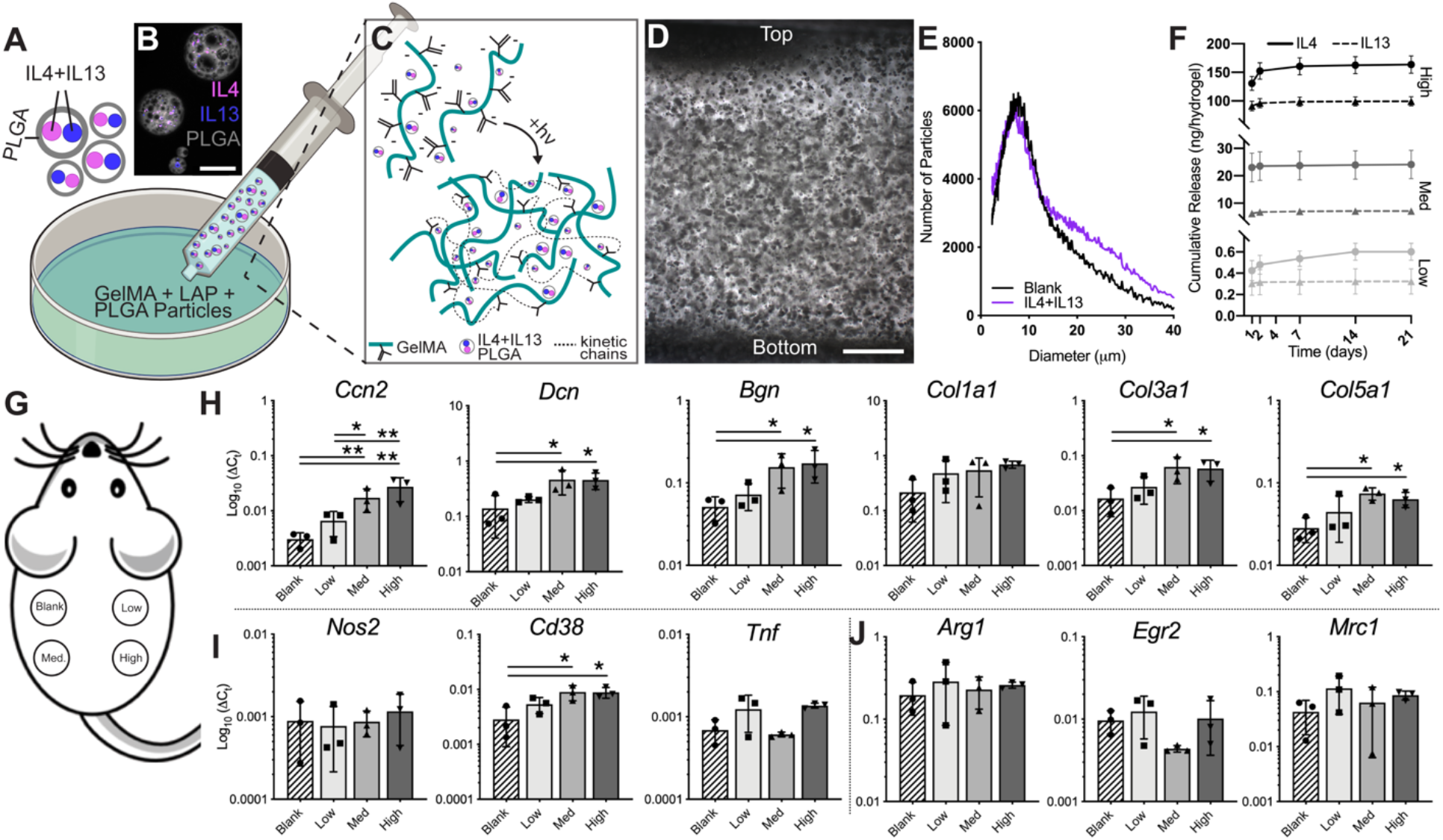
IL4+IL13-loaded PLGA-GelMA hydrogels promote a dose dependent increase in ECM-related gene expression after 21 days in vivo. A) IL4+IL13-loaded PLGA microparticles were fabricated using a double emulsion technique and B) visualized with fluorescently labeled proteins via representative confocal microscopy. Scale bar = 25μm. C) IL4+IL13-loaded PLGA microparticles were embedded in UV-crosslinked GelMA hydrogels. D) Representative image of homogenous PLGA (round black dots) microparticle distribution throughout hydrogel. Scale bar = 500μm. E) Microparticle size distribution; n=3 batches of microparticles with approximately 25,000 particles analyzed in each sample. Data represent mean diameter (Blank: 8.8 ± 6.5μm, IL4+IL13: 9.7 ± 8.0μm). F) Cumulative release profiles of IL4 and IL13 from PLGA-GelMA hydrogels prepared with three different doses (Low, Medium (Med), and High), n=6 experimental replicates. G) Hydrogels were implanted subcutaneously in mice for 21 days. qRTPCR gene expression analysis of H) ECM genes, I) M1 genes, and J) M2 genes. Data represented as mean ± SD, *p<0.05, **p<0.01.

A follow-up study conducted with only the blank and high dose of IL4+IL13 groups suggested that IL4+IL13 released from the high dose group may affect cells surrounding the blank hydrogels within the back of the same mouse, because expression of M2 markers was higher when hydrogels from each group were implanted in the same mouse compared to separate mice (**Suppl. Fig. 10**). Therefore, for all subsequent analyses, only one type of hydrogel was implanted per mouse (two hydrogels per mouse). NanoString multiplex gene expression analysis was used to measure expression of a 70-gene panel representing multiple M1 and M2 markers as well as genes associated with ECM regulation, fibroblast activity, proteases, angiogenesis, and immune signaling pathways. Heatmap visualization of the data and principal component analyses showed differences in gene expression resulting from both time point and inclusion of IL4+IL13 (**Fig. 4A, B**). In addition, potential sex differences were noted (**Fig. 4B**, **Suppl. Fig. 11**), although the small numbers of replicates in each sex precluded statistical analysis of this effect.

**Figure 4.**
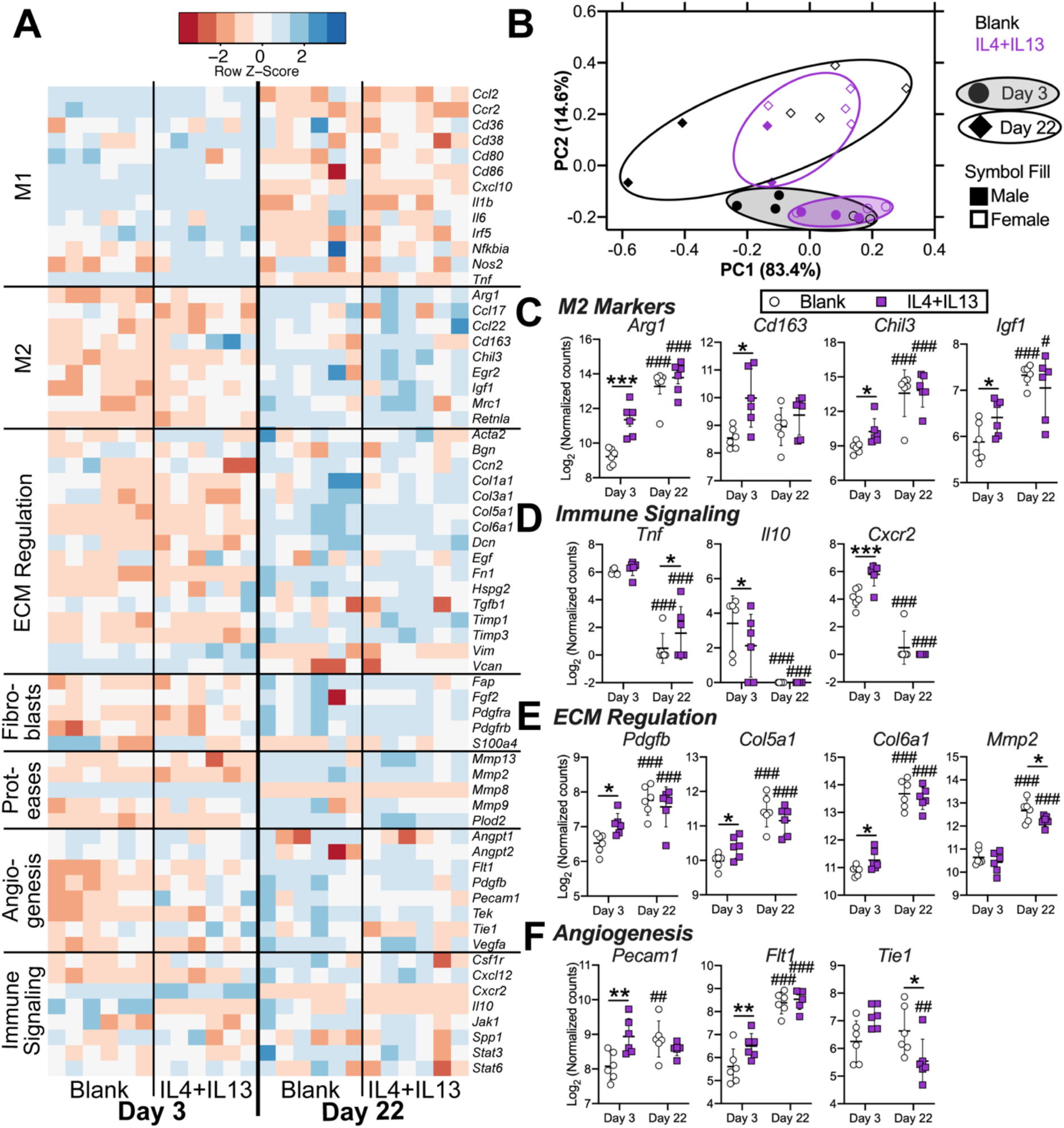
IL4+IL13-releasing hydrogels promote upregulation in genes related to M2 markers, ECM assembly, and angiogenesis in vivo. A) Heatmap of all genes represented as the row Z-score of log-transformed normalized data, B) Principal component analysis, C-F) Differentially expressed genes organized by C) M2 markers, D) Immune signaling, E) ECM regulation, and F) Angiogenesis. *, # p<0.05, **, ## p<0.01, ***, ### p<0.001, where # symbols indicate significance compared to corresponding Day 3 data. All genes are shown in **Suppl. Figs. 12-18**. Genes that were not expressed were: *Cxcl11, Ifna2, Il12a, Il2, Il1a, Mmp7*. Data represented as mean ± SD, n=6 per group, per time point.

In general, gene expression of M1 markers decreased from day 3 to day 22, while expression of M2 markers increased over time (**Fig. 4A, Suppl. Figs. 12-13**). The release of IL4+IL13 caused increased expression of the M2 markers *Arg1, Cd163, Chil3*, and *Igf1* compared to the blank control group at day 3 (**Fig. 4C**), while expression of M1 genes was generally unaffected by release of IL4+IL13, except for *Tnf*, which was significantly increased at day 22 (**Fig. 4D**). Expression of genes involved in regulation of ECM assembly generally increased from day 3 to day 22 (**Suppl. Fig. 15**), with IL4+IL13 resulting in significant increases in *Pdgfb, Col5a1*, and *Col6a1* at day 3 and decreased *Mmp2* at day 22 (**Fig. 4E**). Expression of fibroblast markers *(Fap, Pdgfra*, and *Pdgfrb)* and other protease-encoding genes *(Mmp13* and *Plod2)* also increased from day 3 to day 22, with no significant differences due to IL4+IL13 (**Suppl. Figs. 16 and 17**). Finally, release of IL4+IL13 caused significant increases in the expression of the angiogenesis-related genes *Flt1, Pecam1*, and *Tie1*, at day 3 compared to the blank group (**Fig. 4F, Suppl. Fig. 18**).

### 3.4 Effects of IL4+IL13 release on fibrous capsule properties

Histological and biochemical analyses of the hydrogel-tissue explants from the day 22 time point were used to assess whether release of M2-promoting IL4+IL13 could modulate ECM assembly within the host foreign body response. While release of IL4+IL13 yielded an upward trend in fibrous capsule thickness and total fibrous capsule area (**Fig. 5A-C, Suppl. Figs. 19-20**), second harmonic generation (SHG) microscopy (**Fig. 5D-F**) showed hydrogels releasing IL4+IL13 resulted in fibrous capsules with significantly fewer straight fibers and decreased alignment compared to the blank control (**Fig. 5F**). Integrated density values, which are related to total collagen content, did not differ between the groups (**Suppl. Fig. 21**). Total collagen content and total sGAG content were also increased in fibrous capsules surrounding IL4+IL13-releasing hydrogels, although this effect was only significant for sGAG content (**Fig. 5G, H**).

**Figure 5.**
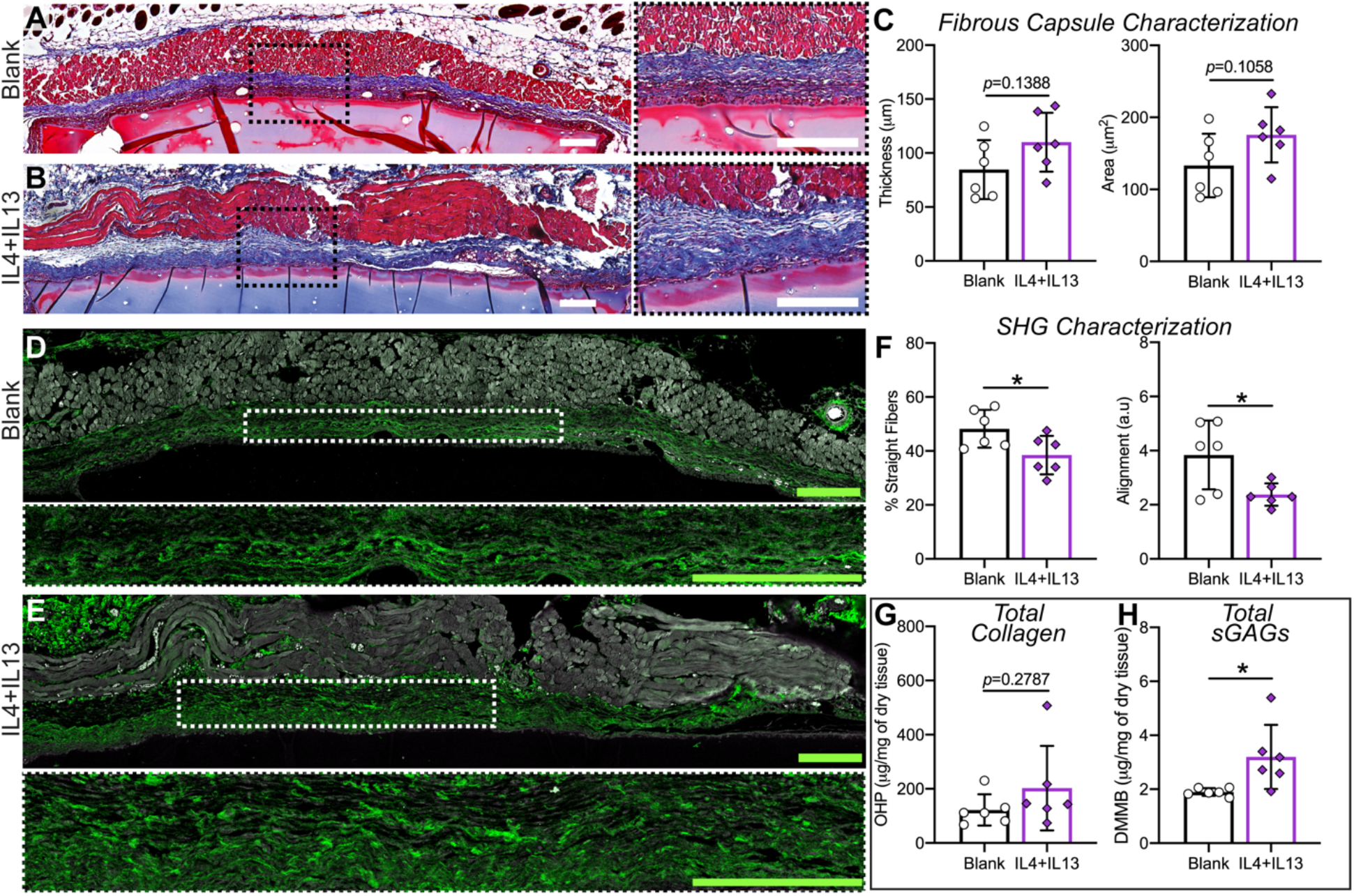
IL4+IL13-releasing hydrogels promote a fibrous capsule with less aligned ECM and increased sulfated glycosaminoglycans (sGAGs). A-B) Representative images of tissue explants (skin-biomaterial interface) of Masson’s Trichrome staining. Scale bar = 100μm. C) Fibrous capsule characterization of thickness and area using image analysis. D-E) Second harmonic generation microscopy was used to analyze fibrillar collagen architecture of the fibrous capsule; regions of interest on each sample were isolated and quantitatively analyzed for F) % straight collagen fibers and fiber alignment. Biochemical analyses were performed on tissue explants for G) total collagen measured using the orthohydroxyproline assay and H) total sGAGs using the DMMB assay. Data represented as mean ± SD, n=6. *p<0.05.

## 4. Discussion

Macrophages exist on a wide spectrum of diverse phenotypes and normally transition from a pro-inflammatory (M1) population to a heterogenous population of M2 macrophages during successful wound healing and tissue repair. Therefore, hybrid M1/M2 phenotypes are likely a normal part of healthy wound healing. On the other hand, fibrosis is also characterized by both M1 and M2 markers, although it is generally considered to be a Th2-driven process regulated primarily by M2-type macrophages. In this study, we aimed to resolve this discrepancy by generating hybrid M1/M2 phenotypes in vitro and directly investigating their effects on fibroblast behavior and ECM assembly *in vitro*, as well as assessing the effects of shifting the population towards M2 on ECM assembly surrounding subcutaneous implanted hydrogels *in vivo*. We found that 1) hybrid M1/M2 macrophage signals induced fibroblast matrix deposition with significantly less aligned matrix compared to M1 or M2 signals alone *in vitro*, and 2) that shifting the balance of hybrid M1/M2 phenotypes that developed around implanted hydrogels *in vivo* towards M2 via delivery of IL4+IL13 increased production of ECM that was also less aligned. Since reduced fiber alignment is associated with less fibrotic tissues [1–4], these data suggest that hybrid M1/M2 macrophages may promote the formation of more easily remodeled ECM, while M2 macrophages enhance ECM deposition. These findings are important because ECM deposition is a requirement for successful wound healing, as long as it can be remodeled appropriately. Biomaterials that affect this balance *in vivo* have potential to reduce implant-associated scarring and improve function.

Numerous studies have shown that macrophage depletion or the controlled release of anti-inflammatory drugs results in significantly reduced fibrous capsule formation [41–44]. However, reports surrounding the effects of macrophage phenotype have been mixed, with some studies reporting an association between higher M2:M1 ratios and thicker fibrous capsules [18, 45, 46] and others reporting the opposite [47]. Outside of fibrous capsule formation, Badylak, Brown, and colleagues have reported improved remodeling outcomes for biomaterials that promoted higher M2:M1 ratios [48, 49]. The present study shows that the combination of M1- and M2-promoting stimuli causes macrophages to take on a unique, hybrid M1/M2 phenotype with potentially beneficial effects on fibroblast behavior, and that increasing the M2:M1 ratio in vivo through the use of IL4+IL13 both increased matrix deposition and made it less aligned.

In addition to promoting reduced fiber alignment and increased sGAG deposition, the delivery of IL4 and IL13 *in vivo* increased the expression of genes related to the regulation of ECM assembly, especially *Col5a1* and *Col6a1*. Collagen VI in particular has been shown to be important for the generation of more randomly aligned ECM [50]. Collagen V plays key roles in regulating the initial fibrillogenesis of collagen I, and its assembly with collagen I suggests a role in reducing alignment, because its hinged, non-helical N-terminal projects outward from the fibril gap region, which inhibits fibril lateral growth and increases interfibril spacing [51]. Studies of model gels prepared with collagen I and collagen V also support a role for collagen V in reducing fibril alignment [52]. Furthermore, collagen V was recently found to play a crucial role in limiting scar size and regulating mechanical properties after cardiac injury [53]. M2 markers that were increased with IL4+IL13 delivery include *Arg1*, which leads to the production of proline, a key amino acid required for collagen deposition [54], and the growth factors *Igf1* and *Pdgfb*, which increase tissue deposition through promotion of fibroblast proliferation [55]. *In vitro*-cultivated M2 macrophages also increased expression of genes encoding perlecan as well as fibronectin. Regulation of cell-ECM interaction and ECM assembly by fibronectin in particular is critical for development of tissue with native-like architecture [56]. One study showed that fibrous capsules surrounding polyethylene terapthalate disks implanted subcutaneously in mice were two-fold thicker in mice genetically depleted of fibronectin, with three times more foreign body giant cells [57]. M2-conditioned media also caused fibroblasts to increase expression of these genes as well as *Col3a1*, although many of these genes were also upregulated in M1- and hybrid M1/M2-conditioned media. All of these proteins are critical regulators of collagen I fibrillogenesis [58]. An important caveat is that only gene expression was measured in this study, and production of the proteins may differ from the results presented here. Nonetheless, these findings support the conclusion that M2 macrophages are likely involved in more complex regulation of ECM assembly beyond a role of simply increasing tissue deposition.

Previous *in vitro* studies of macrophage-fibroblast crosstalk have generally supported a pro-fibrotic role for M2 macrophages, because M2 macrophages promoted fibroblast migration, proliferation, and collagen deposition, while M1 macrophages caused fibroblasts to upregulate matrix-degrading enzymes [48, 59–61]. Our results are in agreement with these studies, in that M2 macrophages caused fibroblasts to upregulate numerous genes associated with cell-ECM interactions and produced a highly aligned matrix. In contrast, the M1-stimulating factors IFNγ and LPS apparently had anti-fibrotic effects on macrophages stimulated with IL4 and IL13, in that they caused fibroblasts to produce matrix with thicker fibers that was less aligned. Similarly, IFNγ has also been shown to inhibit the pro-fibrotic effects of IL4 and IL13 on fibroblasts [62, 63].

Although our *in vitro* studies suggested that M2 macrophages increase fibroblast matrix deposition, it was somewhat surprising that IL4+IL13-releasing hydrogels increased matrix deposition *in vivo* considering two other studies have reported decreased fibrous capsule formation around biomaterials releasing IL4 [21, 22]. Several experimental differences may explain this discrepancy, including the use of different biomaterials, animal species, implantation sites, and time points. A major difference is that our study utilized the release of IL4 together with IL13 in order to enhance the M2 phenotype. IL4 and IL13 are complimentary in their promotion of the M2 phenotype, and are often used in combination to polarize macrophages *in vitro*. While both IL4 and IL13 have been linked to fibrosis, studies that investigated the interactions between these two cytokines in parasite-induced liver fibrosis suggested that IL13 may potentiate fibrotic effects of IL4 [64–66]. Clearly, there is a critical need to further understand how modulation of macrophage phenotype interacts with other biomaterial design parameters as well as microenvironmental cues to affect the foreign body response.

It is important to note that the *in vitro* studies reported here clearly show the effects of different macrophage phenotypes on fibroblast behavior in the absence of IL4 and IL13, since the conditioned media was prepared without these cytokines and macrophages are not major producers of them. However, the effects of IL4+IL13 delivery *in vivo* are likely a combined effect of actions on both macrophages and fibroblasts, although the majority of cells in the fibrous capsule were macrophages, since IL4 and IL13 are both known to have direct, fibrotic effects on both macrophages and fibroblasts [63, 67]. Future studies should thoroughly investigate the effects on each cell type, such as through the use of mice whose myeloid cells but not fibroblasts are devoid of IL4 or IL13 receptors [68].

Another important consideration in interpretation of these results in light of the literature is that there are many distinct phenotypes that have all been described as “M2.” For example, polarization with IL4 and IL13, which we used in this study, is considered to promote an M2a phenotype, while polarization with IL10 is considered to promote an M2c phenotype characterized by distinct surface markers like CD163 and increased secretion of numerous matrix-degrading enzymes like matrix metalloprotease (MMP) −7 and −8 [69], suggesting distinct effects on ECM remodeling. Furthermore, transcriptomic studies of macrophages isolated from the *in vivo* environment suggest that they are difficult to characterize as precisely as macrophage phenotypes studied *in vitro* [9, 19, 70]. Therefore, it appears likely that the choice of particular M2 markers used to characterize macrophages strongly affects interpretation, and that large panels of markers should be used to facilitate comparison between studies.

The results presented here suggest that shifting the balance of macrophage phenotype towards M2 (via IL4 and IL13) influences ECM assembly and microstructure, and motivates the need for more studies investigating the interplay of macrophage phenotype, GAG production, and regulation of ECM assembly and microstructure in the foreign body response and tissue repair.

## Conclusions

In summary, we used *in vitro* studies to understand fibroblast behavior in response to different phenotypes of macrophages, and *in vivo* studies to investigate the effects of increasing M2 activity on formation of the fibrous capsule surrounding biomaterials. We report that hybrid M1/M2 macrophages encourage the production of more native-like (less fibrotic) ECM compared to a predominantly M2 macrophage phenotype, and that increasing M2 activity increases ECM production. These findings have important implications for the design of biomaterials that promote enhanced tissue remodeling and function.

## Acknowledgments

The authors gratefully acknowledge assistance from the Drexel University Cell Imaging Core and the Anatomical Pathology Shared Services of the Sidney Kimmel Medical College, Department of Pathology, Anatomy, & Cell Biology, Jefferson University. Schematics were created with BioRender.com. This work was funded by NHLBI R01 HL130037 (to KLS).

## Data Availability

All processed data are included in main or supplementary figures. Raw data are available from the corresponding author upon request.

## References

[1] P.D. Verhaegen, J.V. Marle, A. Kuehne, H.J. Schouten, E.A. Gaffney, P.K. Maini, E. Middelkoop, P.P. Zuijlen, Collagen bundle morphometry in skin and scar tissue: a novel distance mapping method provides superior measurements compared to Fourier analysis, J Microsc, 245 (2012) 82–89.

[2] P.P. van Zuijlen, J.J. Ruurda, H.A. van Veen, J. van Marle, A.J. van Trier, F. Groenevelt, R.W. Kreis, E. Middelkoop, Collagen morphology in human skin and scar tissue: no adaptations in response to mechanical loading at joints, Burns: journal of the International Society for Burn Injuries, 29 (2003) 423–431.

[3] P.P. van Zuijlen, H.J. de Vries, E.N. Lamme, J.E. Coppens, J. van Marle, R.W. Kreis, E. Middelkoop, Morphometry of dermal collagen orientation by Fourier analysis is superior to multiobserver assessment, The Journal of pathology, 198 (2002) 284–291.

[4] H.J. de Vries, D.N. Enomoto, J. van Marle, P.P. van Zuijlen, J.R. Mekkes, J.D. Bos, Dermal organization in scleroderma: the fast Fourier transform and the laser scatter method objectify fibrosis in nonlesional as well as lesional skin, Laboratory investigation; a journal of technical methods and pathology, 80 (2000) 1281–1289.

[5] L. Arnold, A. Henry, F. Poron, Y. Baba-Amer, N. van Rooijen, A. Plonquet, R.K. Gherardi, B. Chazaud, Inflammatory monocytes recruited after skeletal muscle injury switch into antiinflammatory macrophages to support myogenesis, J Exp Med, 204 (2007) 1057–1069.

[6] P.L. Graney, S. Ben-Shaul, S. Landau, A. Bajpai, J.M. Eager, B. Singh, J.M. Eager, A.R. Cohen, S. Levenberg, K.L. Spiller, Macrophages of diverse phenotypes drive vascularization of engineered tissues, Science Advances, 6 (2020) eaay6391.

[7] K.L. Spiller, T.J. Koh, Macrophage-based therapeutic strategies in regenerative medicine, Advanced drug delivery reviews, (2017).

[8] E.M. O’Brien, G.E. Risser, K.L. Spiller, Sequential drug delivery to modulate macrophage behavior and enhance implant integration, Advanced drug delivery reviews, (2019).

[9] S.D. Sommerfeld, C. Cherry, R.M. Schwab, L. Chung, D.R. Maestas, Jr., P. Laffont, J.E. Stein, A. Tam, S. Ganguly, F. Housseau, J.M. Taube, D.M. Pardoll, P. Cahan, J.H. Elisseeff, Interleukin-36gamma-producing macrophages drive IL-17-mediated fibrosis, Sci Immunol, 4 (2019).

[10] T.A. Wynn, L. Barron, Macrophages: master regulators of inflammation and fibrosis, Semin Liver Dis, 30 (2010) 245–257.

[11] K. Zhang, M. Gharaee-Kermani, B. McGarry, D. Remick, S.H. Phan, TNF-alpha-mediated lung cytokine networking and eosinophil recruitment in pulmonary fibrosis, Journal of immunology, 158 (1997) 954–959.

[12] W. Liu, I. Ding, K. Chen, J. Olschowka, J. Xu, D. Hu, G.R. Morrow, P. Okunieff, Interleukin 1beta (IL1B) signaling is a critical component of radiation-induced skin fibrosis, Radiat Res, 165 (2006) 181–191.

[13] F. Ma, Y. Li, L. Jia, Y. Han, J. Cheng, H. Li, Y. Qi, J. Du, Macrophage-stimulated cardiac fibroblast production of IL-6 is essential for TGF beta/Smad activation and cardiac fibrosis induced by angiotensin II, PloS one, 7 (2012) e35144.

[14] W.A. Border, N.A. Noble, Transforming growth factor beta in tissue fibrosis, N Engl J Med, 331 (1994) 1286–1292.

[15] E.S. Yi, H. Lee, S. Yin, P. Piguet, I. Sarosi, S. Kaufmann, J. Tarpley, N.S. Wang, T.R. Ulich, Platelet-derived growth factor causes pulmonary cell proliferation and collagen deposition in vivo, The American journal of pathology, 149 (1996) 539–548.

[16] T. Inoue, S. Fujishima, E. Ikeda, O. Yoshie, N. Tsukamoto, S. Aiso, N. Aikawa, A. Kubo, K. Matsushima, K. Yamaguchi, CCL22 and CCL17 in rat radiation pneumonitis and in human idiopathic pulmonary fibrosis, Eur Respir J, 24 (2004) 49–56.

[17] K.L. Spiller, R.R. Anfang, K.J. Spiller, J. Ng, K.R. Nakazawa, J.W. Daulton, G. Vunjak-Novakovic, The role of macrophage phenotype in vascularization of tissue engineering scaffolds, Biomaterials, 35 (2014) 4477–4488.

[18] T. Yu, W. Wang, S. Nassiri, T. Kwan, C. Dang, W. Liu, K.L. Spiller, Temporal and spatial distribution of macrophage phenotype markers in the foreign body response to glutaraldehyde-crosslinked gelatin hydrogels, Journal of biomaterials science. Polymer edition, 27 (2016) 721–742.

[19] J.E. Mooney, K.M. Summers, M. Gongora, S.M. Grimmond, J.H. Campbell, D.A. Hume, B.E. Rolfe, Transcriptional switching in macrophages associated with the peritoneal foreign body response, Immunol Cell Biol, 92 (2014) 518–526.

[20] W.J. Kao, A.K. McNally, A. Hiltner, J.M. Anderson, Role for interleukin-4 in foreign-body giant cell formation on a poly(etherurethane urea) in vivo, Journal of biomedical materials research, 29 (1995) 1267–1275.

[21] D. Hachim, S.T. LoPresti, C.C. Yates, B.N. Brown, Shifts in macrophage phenotype at the biomaterial interface via IL-4 eluting coatings are associated with improved implant integration, Biomaterials, 112 (2017) 95–107.

[22] R.P. Tan, A.H.P. Chan, S. Wei, M. Santos, B.S.L. Lee, E.C. Filipe, B. Akhavan, M.M. Bilek, M.K.C. Ng, Y. Xiao, S.G. Wise, Bioactive Materials Facilitating Targeted Local Modulation of Inflammation, JACC Basic Transl Sci, 4 (2019) 56–71.

[23] E.M. O’Brien, K.L. Spiller, Prior pro-inflammatory polarization changes the subsequent response of macrophages to IL4, In revisions at J Leukocyte Biology, (2020).

[24] N.A. Hotaling, K. Bharti, H. Kriel, C.G. Simon, DiameterJ: A validated open source nanofiber diameter measurement tool, Biomaterials, 61 (2015) 327–338.

[25] Q. Li, F. Qu, B. Han, C. Wang, H. Li, R.L. Mauck, L. Han, Micromechanical anisotropy and heterogeneity of the meniscus extracellular matrix, Acta biomaterialia, 54 (2017) 356–366.

[26] C.E. Witherel, P.L. Graney, D.O. Freytes, M.S. Weingarten, K.L. Spiller, Response of human macrophages to wound matrices in vitro, Wound Repair Regen, 24 (2016) 514–524.

[27] J.L. Hutter, J. Bechhoefer, Calibration of atomic-force microscope tips, Review of Scientific Instruments, 64 (1993) 1868–1873.

[28] L. Han, E.H. Frank, J.J. Greene, H.-Y. Lee, H.-H.K. Hung, A.J. Grodzinsky, C. Ortiz, Time-dependent nanomechanics of cartilage, Biophys J, 100 (2011) 1846–1854.

[29] E.K. Dimitriadis, F. Horkay, J. Maresca, B. Kachar, R.S. Chadwick, Determination of elastic moduli of thin layers of soft material using the atomic force microscope, Biophys J, 82 (2002) 2798–2810.

[30] I.G. Kim, C.H. Gil, J. Seo, S.J. Park, R. Subbiah, T.H. Jung, J.S. Kim, Y.H. Jeong, H.M. Chung, J.H. Lee, M.R. Lee, S.H. Moon, K. Park, Mechanotransduction of human pluripotent stem cells cultivated on tunable cell-derived extracellular matrix, Biomaterials, 150 (2018) 100–111.

[31] P.H. Mott, J.R. Dorgan, C.M. Roland, The bulk modulus and Poisson’s ratio of “incompressible” materials, Journal of Sound and Vibration, 312 (2008) 572–575.

[32] S. Preibisch, S. Saalfeld, P. Tomancak, Globally optimal stitching of tiled 3D microscopic image acquisitions, Bioinformatics, 25 (2009) 1463–1465.

[33] T.J.A. Mommersteeg, J.M.G. Kauer, R. Huiskes, L. Blankevoort, Method to determine collagen density distributions in fibrous tissues, Journal of Orthopaedic Research, 11 (1993) 612–616.

[34] V.J. Coulson-Thomas, t. ferreira, Dimethylmethylene Blue Assay (DMMB), Bio-protocol, 4 (2014) e1236.

[35] A. Case, B.K. Brisson, A.C. Durham, S. Rosen, J. Monslow, E. Buza, P. Salah, J. Gillem, G. Ruthel, S. Veluvolu, V. Kristiansen, E. Pure, D.C. Brown, K.U. Sorenmo, S.W. Volk, Identification of prognostic collagen signatures and potential therapeutic stromal targets in canine mammary gland carcinoma, PloS one, 12 (2017) e0180448.

[36] S. Rosen, B.K. Brisson, A.C. Durham, C.M. Munroe, C.J. McNeill, D. Stefanovski, K.U. Sorenmo, S.W. Volk, Intratumoral collagen signatures predict clinical outcomes in feline mammary carcinoma, PloS one, 15 (2020) e0236516.

[37] J.S. Bredfeldt, Y. Liu, M.W. Conklin, P.J. Keely, T.R. Mackie, K.W. Eliceiri, Automated quantification of aligned collagen for human breast carcinoma prognosis, J Pathol Inform, 5 (2014) 28.

[38] B.K. Brisson, E.A. Mauldin, W. Lei, L.K. Vogel, A.M. Power, A. Lo, D. Dopkin, C. Khanna, R.G. Wells, E. Pure, S.W. Volk, Type III Collagen Directs Stromal Organization and Limits Metastasis in a Murine Model of Breast Cancer, The American journal of pathology, 185 (2015) 1471–1486.

[39] Y. Benjamini, A.M. Krieger, D. Yekutieli, Adaptive linear step-up procedures that control the false discovery rate, Biometrika, 93 (2006) 491–507.

[40] M.E. Glickman, S.R. Rao, M.R. Schultz, False discovery rate control is a recommended alternative to Bonferroni-type adjustments in health studies, J Clin Epidemiol, 67 (2014) 850–857.

[41] N.M. Vacanti, H. Cheng, P.S. Hill, J.D. Guerreiro, T.T. Dang, M. Ma, S. Watson, N.S. Hwang, R. Langer, D.G. Anderson, Localized delivery of dexamethasone from electrospun fibers reduces the foreign body response, Biomacromolecules, 13 (2012) 3031–3038.

[42] E. Blanco, B.D. Weinberg, N.T. Stowe, J.M. Anderson, J. Gao, Local release of dexamethasone from polymer millirods effectively prevents fibrosis after radiofrequency ablation, Journal of biomedical materials research. Part A, 76 (2006) 174–182.

[43] J.E. Mooney, B.E. Rolfe, G.W. Osborne, D.P. Sester, N. van Rooijen, G.R. Campbell, D.A. Hume, J.H. Campbell, Cellular plasticity of inflammatory myeloid cells in the peritoneal foreign body response, The American journal of pathology, 176 (2010) 369–380.

[44] E. Dondossola, B.M. Holzapfel, S. Alexander, S. Filippini, D.W. Hutmacher, P. Friedl, Examination of the foreign body response to biomaterials by nonlinear intravital microscopy, Nat Biomed Eng, 1 (2016).

[45] E.M. Sussman, M.C. Halpin, J. Muster, R.T. Moon, B.D. Ratner, Porous implants modulate healing and induce shifts in local macrophage polarization in the foreign body reaction, Annals of biomedical engineering, 42 (2014) 1508–1516.

[46] S.M. van Putten, D.T. Ploeger, E.R. Popa, R.A. Bank, Macrophage phenotypes in the collagen-induced foreign body reaction in rats, Acta biomaterialia, 9 (2013) 6502–6510.

[47] L. Zhang, Z. Cao, T. Bai, L. Carr, J.R. Ella-Menye, C. Irvin, B.D. Ratner, S. Jiang, Zwitterionic hydrogels implanted in mice resist the foreign-body reaction, Nature biotechnology, 31 (2013) 553–556.

[48] B.N. Brown, R. Londono, S. Tottey, L. Zhang, K.A. Kukla, M.T. Wolf, K.A. Daly, J.E. Reing, S.F. Badylak, Macrophage phenotype as a predictor of constructive remodeling following the implantation of biologically derived surgical mesh materials, Acta biomaterialia, 8 (2012) 978–987.

[49] S.F. Badylak, J.E. Valentin, A.K. Ravindra, G.P. McCabe, A.M. Stewart-Akers, Macrophage phenotype as a determinant of biologic scaffold remodeling, Tissue engineering. Part A, 14 (2008) 1835–1842.

[50] G. Theocharidis, Z. Drymoussi, A.P. Kao, A.H. Barber, D.A. Lee, K.M. Braun, J.T. Connelly, Type VI Collagen Regulates Dermal Matrix Assembly and Fibroblast Motility, The Journal of investigative dermatology, 136 (2016) 74–83.

[51] T.F. Linsenmayer, E. Gibney, F. Igoe, M.K. Gordon, J.M. Fitch, L.I. Fessler, D.E. Birk, Type V collagen: molecular structure and fibrillar organization of the chicken alpha 1(V) NH2-terminal domain, a putative regulator of corneal fibrillogenesis, The Journal of cell biology, 121 (1993) 1181–1189.

[52] V. Ajeti, O. Nadiarnykh, S.M. Ponik, P.J. Keely, K.W. Eliceiri, P.J. Campagnola, Structural changes in mixed Col I/Col V collagen gels probed by SHG microscopy: implications for probing stromal alterations in human breast cancer, Biomed Opt Express, 2 (2011) 2307–2316.

[53] T. Yokota, J. McCourt, F. Ma, S. Ren, S. Li, T.H. Kim, Y.Z. Kurmangaliyev, R. Nasiri, S. Ahadian, T. Nguyen, X.H.M. Tan, Y. Zhou, R. Wu, A. Rodriguez, W. Cohn, Y. Wang, J. Whitelegge, S. Ryazantsev, A. Khademhosseini, M.A. Teitell, P.Y. Chiou, D.E. Birk, A.C. Rowat, R.H. Crosbie, M. Pellegrini, M. Seldin, A.J. Lusis, A. Deb, Type V Collagen in Scar Tissue Regulates the Size of Scar after Heart Injury, Cell, 182 (2020) 545–562 e523.

[54] M.A. Dunn, M. Rojkind, K.S. Warren, P.K. Hait, L. Rifas, S. Seifter, Liver collagen synthesis in murine schistosomiasis, The Journal of clinical investigation, 59 (1977) 666–674.

[55] M. Hetzel, M. Bachem, D. Anders, G. Trischler, M. Faehling, Different effects of growth factors on proliferation and matrix production of normal and fibrotic human lung fibroblasts, Lung, 183 (2005) 225–237.

[56] I. Wierzbicka-Patynowski, J.E. Schwarzbauer, The ins and outs of fibronectin matrix assembly, J Cell Sci, 116 (2003) 3269–3276.

[57] B.G. Keselowsky, A.W. Bridges, K.L. Burns, C.C. Tate, J.E. Babensee, M.C. LaPlaca, A.J. Garcia, Role of plasma fibronectin in the foreign body response to biomaterials, Biomaterials, 28 (2007) 3626–3631.

[58] K.E. Kadler, A. Hill, E.G. Canty-Laird, Collagen fibrillogenesis: fibronectin, integrins, and minor collagens as organizers and nucleators, Curr Opin Cell Biol, 20 (2008) 495–501.

[59] D.T. Ploeger, N.A. Hosper, M. Schipper, J.A. Koerts, S. de Rond, R.A. Bank, Cell plasticity in wound healing: paracrine factors of M1/ M2 polarized macrophages influence the phenotypical state of dermal fibroblasts, Cell communication and signaling: CCS, 11 (2013) 29.

[60] Z. Zhu, J. Ding, Z. Ma, T. Iwashina, E.E. Tredget, Alternatively activated macrophages derived from THP-1 cells promote the fibrogenic activities of human dermal fibroblasts, Wound repair and regeneration: official publication of the Wound Healing Society [and] the European Tissue Repair Society, 25 (2017) 377–388.

[61] M.R. Fernando, M.A. Giembycz, D.M. McKay, Bidirectional crosstalk via IL-6, PGE2 and PGD2 between murine myofibroblasts and alternatively activated macrophages enhances antiinflammatory phenotype in both cells, Br J Pharmacol, 173 (2016) 899–912.

[62] G.D. Sempowski, S. Derdak, R.P. Phipps, Interleukin-4 and interferon-gamma discordantly regulate collagen biosynthesis by functionally distinct lung fibroblast subsets, Journal of cellular physiology, 167 (1996) 290–296.

[63] A. Saito, H. Okazaki, I. Sugawara, K. Yamamoto, H. Takizawa, Potential action of IL-4 and IL-13 as fibrogenic factors on lung fibroblasts in vitro, Int Arch Allergy Immunol, 132 (2003) 168–176.

[64] P.G. Fallon, E.J. Richardson, G.J. McKenzie, A.N. McKenzie, Schistosome infection of transgenic mice defines distinct and contrasting pathogenic roles for IL-4 and IL-13: IL-13 is a profibrotic agent, Journal of immunology, 164 (2000) 2585–2591.

[65] M.G. Chiaramonte, D.D. Donaldson, A.W. Cheever, T.A. Wynn, An IL-13 inhibitor blocks the development of hepatic fibrosis during a T-helper type 2-dominated inflammatory response, The Journal of clinical investigation, 104 (1999) 777–785.

[66] M.G. Chiaramonte, L.R. Schopf, T.Y. Neben, A.W. Cheever, D.D. Donaldson, T.A. Wynn, IL-13 is a key regulatory cytokine for Th2 cell-mediated pulmonary granuloma formation and IgE responses induced by Schistosoma mansoni eggs, Journal of immunology, 162 (1999) 920–930.

[67] C. Doucet, D. Brouty-Boye, C. Pottin-Clemenceau, G.W. Canonica, C. Jasmin, B. Azzarone, Interleukin (IL) 4 and IL-13 act on human lung fibroblasts. Implication in asthma, The Journal of clinical investigation, 101 (1998) 2129–2139.

[68] J.A. Knipper, S. Willenborg, J. Brinckmann, W. Bloch, T. Maab, R. Wagener, T. Krieg, T. Sutherland, A. Munitz, M.E. Rothenberg, A. Niehoff, R. Richardson, M. Hammerschmidt, J.E. Allen, S.A. Eming, Interleukin-4 Receptor α Signaling in Myeloid Cells Controls Collagen Fibril Assembly in Skin Repair, Immunity, 43 (2015) 803–816.

[69] E.B. Lurier, D. Dalton, W. Dampier, P. Raman, S. Nassiri, N.M. Ferraro, R. Rajagopalan, M. Sarmady, K.L. Spiller, Transcriptome analysis of IL-10-stimulated (M2c) macrophages by nextgeneration sequencing, Immunobiology, 222 (2017) 847–856.

[70] J. Xue, S.V. Schmidt, J. Sander, A. Draffehn, W. Krebs, I. Quester, D. De Nardo, T.D. Gohel, M. Emde, L. Schmidleithner, H. Ganesan, A. Nino-Castro, M.R. Mallmann, L. Labzin, H. Theis, M. Kraut, M. Beyer, E. Latz, T.C. Freeman, T. Ulas, J.L. Schultze, Transcriptome-based network analysis reveals a spectrum model of human macrophage activation, Immunity, 40 (2014) 274–288.

