## SupplementaryFile for "Regulation of extracellular matrix assembly and structure by hybrid M1/M2 macrophages"

| **Supplementary Table 1:** qRT-PCR murine primers | | |
|---|---|---|
| **Gene** | **Forward** | **Reverse** |
| *Acta2* | GACAGCTACGTGGGTGACGAA | TTTTCCATGTCGTCCCAGTTG |
| *Arg1* | CTCCAAGCCAAAGTCCTTAGAG | GGAGCTGTCATTAGGGACATCA |
| *b-actin* | TCACCCACACTGTGCCCATCTACGA | GGATGCCACAGGATTCCATACCCA |
| *Bgn* | CTACGCCCTGGTCTTGGTAA | ACTTTGCGGATACGGTTGTC |
| *Ccl17* | CAAGCTCATCTGTGCAGACC | CGCCTGTAGTGCATAAGAGTCC |
| *Ccl22* | AAGACAGTATCTGCTGCCAGG | GATCGGCACAGATATCTCGG |
| *Ccn2* | CCTGGTCCAGACCACAGAGT | TTTTCCTCCAGGTCAGCTTC |
| *Cd38* | TTGCAAGGGTTCTTGGAAAC | CGCTGCCTCATCTACACTCA |
| *Chil3* | CACCATGGCCAAGCTCATTCTTGT | TATTGGCCTGTCCTTAGCCCAACT |
| *Col1a1* | TTCTCCTGGCAAAGACGGACTCAA | AGGAAGCTGAAGTCATAACCGCCA |
| *Col2a1* | GCTGGTGCACAAGGTCCTAT | ACCCTGCAGTCCAGTGAAAC |
| *Col3a1* | CAGGTCCTAGAGGAAACAGA | TCACCTCCAACTCCAACAATG |
| *Col5a1* | AAGCGTGGGAAACTGCTCTCCTAT | AGCAGTTGTAGGTGACGTTCTGGT |
| *Col6a3* | TCTTGAACGTGTGGCTAACC | CTTTTCTCCAGAAGAACCAGG |
| *Dcn* | TGAGCTTCAACAGCATCACC | AAGTCATTTTGCCCAACTGC |
| *Egr2* | TTGACCAGATGAACGGAGTG | AGCTACTCGGATACGGGAGA |
| *Fn1* | CGAGGTGACAGAGACCACAA | CTGGAGTCAAGCCAGACACA |
| *Il1b* | GCCTTGGGCCTCAAAGGAAAGAATC | GGAAGACACAGATTCCATGGTGAAG |
| *Lum* | TCGAGCTTGATCTCTCCTAT | TGGTCCCAGGATCTTACAGAA |
| *Mrc1* | GGAAAAATGGATCCACACCTTGC | TCTCTTCCTCCACCACCATGCAG |
| *Nos2* | GGAGTGACGGCAAACATGACT | TCGATGCACAACTGGGTGAAC |
| *Retnla* | TCCCAGTGAATACTGATGAGA | CCACTCTGGATCTCCCAAGA |
| *Tnf* | CAGGCGGTGCCTATGTCTC | CGATCACCCCGAAGTTCAGTAG |

| **Supplementary Table 2**: Human NanoString Gene Panel | | |
|---|---|---|
| **Gene** | **Target Identifier** | **Annotation** |
| ACTA2 | NM_001613.1 | ECM |
| AGGF1 | NM_018046.3 | Angiogenesis |
| ANG | NM_001145.4 | M2 |
| BGN | NM_001711.3 | ECM |
| BTG1 | NM_001731.2 | Immune Signaling |
| CABLES1 | NM_001100619.2 | M2 |
| CCL15 | NM_032965.4 | Angiogenesis |
| CCL17 | NM_002987.2 | M2 |
| CCL18 | NM_002988.2 | M2 |
| CCL2 | NM_002982.3 | M1 |

| | | |
|---|---|---|
| CCL22 | NM_002990.3 | M2 |
| CCL24 | NM_002991.2 | M2 |
| CCL26 | NM_006072.4 | M2 |
| CCL5 | NM_002985.2 | Angiogenesis |
| CCL8 | NM_005623.2 | M1 |
| CCR7 | NM_001838.2 | M1 |
| CD163 | NM_004244.4 | M2c |
| CD200R1 | NM_138806.3 | M2 |
| CD80 | NM_005191.3 | M1 |
| CLEC10A | NM_182906.2 | M2 |
| COL1A1 | NM_000088.3 | ECM |
| COL3A1 | NM_000090.3 | ECM |
| COL5A1 | NM_000093.3 | ECM |
| CTGF | NM_001901.2 | ECM |
| CTNNB1 | NM_001098210.1 | Angiogenesis |
| CXCL12 | NM_199168.3 | M1 |
| CXCR4 | NM_003467.2 | Immune Signaling |
| DACT1 | NM_001079520.1 | M2 |
| DCN | NM_001920.3 | ECM |
| EGFL7 | NM_016215.3 | Angiogenesis |
| ETS1 | NM_005238.3 | Angiogenesis |
| FGF2 | NM_002006.4 | ECM |
| FLT1 | NM_002019.4 | Angiogenesis |
| FN1 | NM_212482.1 | ECM |
| FOXO1 | NM_002015.3 | Angiogenesis |
| FOXO3 | NM_001455.2 | Angiogenesis |
| FOXO4 | NM_001170931.1 | Angiogenesis |
| FST | NM_006350.2 | Angiogenesis |
| FYN | NM_002037.3 | Immune Signaling |
| HSPG2 | NM_005529.5 | ECM |
| IDO1 | NM_002164.5 | M1 |
| IGF1 | NM_000618.3 | M2 |
| IL1B | NM_000576.2 | M1 |
| IL6 | NM_000600.3 | M1 |
| JAG1 | NM_000214.2 | Immune Signaling |
| LUM | NM_002345.3 | ECM |
| MARCO | NM_006770.3 | M2c |
| MMP2 | NM_004530.2 | M1 |
| MMP9 | NM_004994.2 | M2c |

| | | |
|---|---|---|
| MRC1 | NM_002438.2 | M2 |
| PDGFA | NM_002607.5 | Angiogenesis |
| PDGFB | NM_033016.2 | Angiogenesis |
| PDGFC | NM_016205.2 | Angiogenesis |
| PDGFRA | NM_006206.3 | ECM |
| PDGFRB | NM_002609.3 | ECM |
| RAMP1 | NM_005855.2 | M2 |
| SPP1 | NM_000582.2 | Immune Signaling |
| STAT3 | NM_003150.3 | Immune Signaling |
| STAT6 | NM_003153.3 | Immune Signaling |
| TGFB1 | NM_000660.3 | ECM |
| TIE1 | NM_005424.2 | Angiogenesis |
| TIMP3 | NM_000362.4 | Immune Signaling |
| TNF | NM_000594.2 | M1 |
| TNFRSF11A | NM_003839.3 | M2 |
| VCAN | NM_004385.3 | ECM |
| VEGFA | NM_001025366.1 | Angiogenesis |
| VEGFB | NM_003377.3 | Angiogenesis |
| VEGFC | NM_005429.2 | Angiogenesis |
| VIM | NM_003380.2 | ECM |
| WNT5A | NM_003392.3 | Immune Signaling |
| GAPDH | NM_001256799.1 | Housekeeping |
| TBP | NM_001172085.1 | Housekeeping |

| **Supplementary Table 3:** Murine NanoString Gene Panel | | |
|---|---|---|
| **Gene** | **Target Identifier** | **Annotation** |
| *Acta2* | NM_007392.2 | ECM |
| *Angpt1* | NM_009640.3 | Angiogenesis |
| *Angpt2* | NM_007426.3 | Angiogenesis |
| *Arg1* | NM_007482.3 | M2 |
| *Bbgn* | NM_007542.4 | ECM |
| *Ccl17* | NM_011332.2 | M2 |
| *Ccl2* | NM_011333.3 | M1 |
| *Ccl22* | NM_009137.2 | M2 |
| *Ccn2* | NM_010217.2 | ECM |
| *Ccr2* | NM_009915.2 | M1 |
| *Cd163* | NM_053094.2 | M2c |
| *Cd36* | NM_007643.3 | M1 |

| | | |
|---|---|---|
| *Cd38* | NM_007646.4 | M1 |
| *Cd80* | NM_009855.2 | M1 |
| *Cd86* | NM_019388.3 | M1 |
| *Chil3* | NM_009892.2 | M2 |
| *Col1a1* | NM_007742.3 | ECM |
| *Col3a1* | NM_009930.1 | ECM |
| *Col5a1* | NM_015734.2 | ECM |
| *Col6a1* | NM_009933.2 | ECM |
| *Csf1r* | NM_001037859.1 | Immune signaling |
| *Cxcl10* | NM_021274.1 | M1 |
| *Cxcl11* | NM_019494.1 | M1 |
| *Cxcl12* | NM_021704.3 | Immune signaling |
| *Cxcr2* | NM_009909.3 | Immune signaling |
| *Dcn* | NM_001190451.1 | ECM |
| *Egf* | NM_010113.3 | ECM |
| *Egr2* | NM_010118.2 | M2 |
| *Fap* | NM_007986.2 | Fibroblasts |
| *Fgf2* | NM_008006.2 | Fibroblasts |
| *Flt1* | NM_010228.3 | Angiogenesis |
| *Fn1* | NM_010233.1 | ECM |
| *Hspg2* | NM_008305.3 | ECM |
| *Ifna* | NM_010503.2 | Immune signaling |
| *Igf1* | NM_001111274.1 | M2 |
| *Il10* | NM_010548.1 | M2c |
| *Il12a* | NM_008351.1 | Immune signaling |
| *Il1a* | NM_010554.4 | M1 |
| *Il1b* | NM_008361.3 | M1 |
| *Il2* | NM_008366.2 | Immune signaling |
| *Il6* | NM_031168.1 | M1 |
| *Irf5* | NM_001252382.1 | M1 |
| *Jak1* | NM_146145.2 | Immune signaling |
| *Mmp13* | NM_008607.1 | Protease |
| *Mmp2* | NM_008610.2 | Protease |
| *Mmp7* | NM_010810.4 | Protease |
| *Mmp8* | NM_008611.4 | Protease |
| *Mmp9* | NM_013599.2 | Protease |
| *Mrc1* | NM_008625.1 | M2 |
| *Nfkbia* | NM_010907.2 | M1 |
| *Nos2* | NM_010927.3 | M1 |

| | | |
|---|---|---|
| *Pdgfb* | NM_011057.3 | Angiogenesis |
| *Pdgfra* | NM_011058.2 | Fibroblasts |
| *Pdgfrb* | NM_008809.1 | Fibroblasts |
| *Pecam1* | NM_008816.2 | Angiogenesis |
| *Plod2* | NM_001142916.1 | Protease |
| *Retnla* | NM_020509.3 | M2 |
| *S100a4* | NM_011311.2 | Fibroblasts |
| *Spp1* | NM_009263.3 | ECM |
| *Stat3* | NM_213659.2 | Immune signaling |
| *Stat6* | NM_009284.2 | Immune signaling |
| *Tek* | NM_013690.2 | Angiogenesis |
| *Tgfb1* | NM_011577.1 | ECM |
| *Tie1* | NM_011587.2 | Angiogenesis |
| *Timp1* | NM_011593.2 | ECM |
| *Timp3* | NM_011595.2 | ECM |
| *Tnf* | NM_013693.2 | M1 |
| *Vcan* | NM_001081249.1 | ECM |
| *Vegfa* | NM_001025250.3 | Angiogenesis |
| *Vim* | NM_011701.4 | ECM |
| *Gusb* | NM_010368.1 | Housekeeping |
| *Tbp* | NM_013684.3 | Housekeeping |

**Supplementary Table 4**: Cumulative release data of Low, Medium, and High dose hydrogels

LOW Dose Hydrogel - Cumulative Release

| Day | Sample 1 | Sample 2 | Sample 3 | Sample 4 | Mean (pg) | St Dev (pg) | Mean (ng) | St Dev (ng) |
|---|---|---|---|---|---|---|---|---|
| 1 | 195.946108 | 194.909244 | 157.158225 | 128.838492 | 169.213017 | 32.4056299 | 0.16921302 | 0.03240563 |
| 2 | 675.442262 | 403.626204 | 400.51704 | 220.695849 | 425.070338 | 187.543548 | 0.42507034 | 0.18754355 |
| 4 | 725.937896 | 403.626204 | 472.913049 | 314.785893 | 479.315761 | 176.694141 | 0.47931576 | 0.17669414 |
| 7 | 725.937896 | 432.805779 | 573.357181 | 416.834181 | 537.233759 | 144.124269 | 0.53723376 | 0.14412427 |
| 14 | 801.925598 | 563.343602 | 624.115372 | 416.834181 | 601.554688 | 159.414804 | 0.60155469 | 0.1594148 |
| 21 | 801.925598 | 563.343602 | 624.115372 | 416.834181 | 601.554688 | 159.414804 | 0.60155469 | 0.1594148 |

MEDIUM Dose Hydrogel - Cumulative Release

| Day | Sample 1 | Sample 2 | Sample 3 | Sample 4 | Mean (pg) | St Dev (pg) | Mean (ng) | St Dev (ng) |
|---|---|---|---|---|---|---|---|---|
| 1 | 18087.7363 | 7066.136 | 22637.826 | 31551.766 | 19835.8661 | 10185.5331 | 19.8358661 | 10.1855331 |
| 2 | 20653.6427 | 9322.90165 | 29130.1649 | 32887.4489 | 22998.5395 | 10454.807 | 22.9985395 | 10.454807 |
| 4 | 21127.7724 | 9840.71315 | 29674.5492 | 33400.7125 | 23510.9368 | 10461.8715 | 23.5109368 | 10.4618715 |
| 7 | 21320.3842 | 10107.9625 | 30012.9682 | 33442.6531 | 23720.992 | 10411.1663 | 23.720992 | 10.4111663 |

| | | | | | | | | |
|---|---|---|---|---|---|---|---|---|
| 14 | 21756.518 | 10341.3255 | 30244.1324 | 33442.6531 | 23946.1573 | 10323.6604 | 23.9461573 | 10.3236604 |
| 21 | 21900.689 | 10428.1974 | 30549.1791 | 33655.8859 | 24133.4878 | 10402.8447 | 24.1334878 | 10.4028447 |
| HIGH Dose Hydrogel - Cumulative Release | | | | | | | | |
| Day | Sample 1 | Sample 2 | Sample 3 | Sample 4 | Mean (pg) | St Dev (pg) | Mean (ng) | St Dev (ng) |
| 1 | 86257.0062 | 124197.124 | 77422.7857 | 118050.819 | 101481.934 | 23102.3006 | 101.481934 | 23.1023006 |
| 2 | 110458.615 | 147909.715 | 109735.361 | 154799.4 | 130725.773 | 23987.3746 | 130.725773 | 23.9873746 |
| 4 | 122808.262 | 182672.342 | 133793.093 | 169785.358 | 152264.764 | 28521.8054 | 152.264764 | 28.5218054 |
| 7 | 132232.487 | 195619.99 | 139929.416 | 174781.233 | 160640.782 | 29774.0897 | 160.640782 | 29.7740897 |
| 14 | 134197.464 | 197722.336 | 141498.223 | 176236.994 | 162413.754 | 29840.668 | 162.413754 | 29.840668 |
| 21 | 135399.359 | 199365.094 | 142421.455 | 177167 | 163588.227 | 30038.9293 | 163.588227 | 30.0389293 |

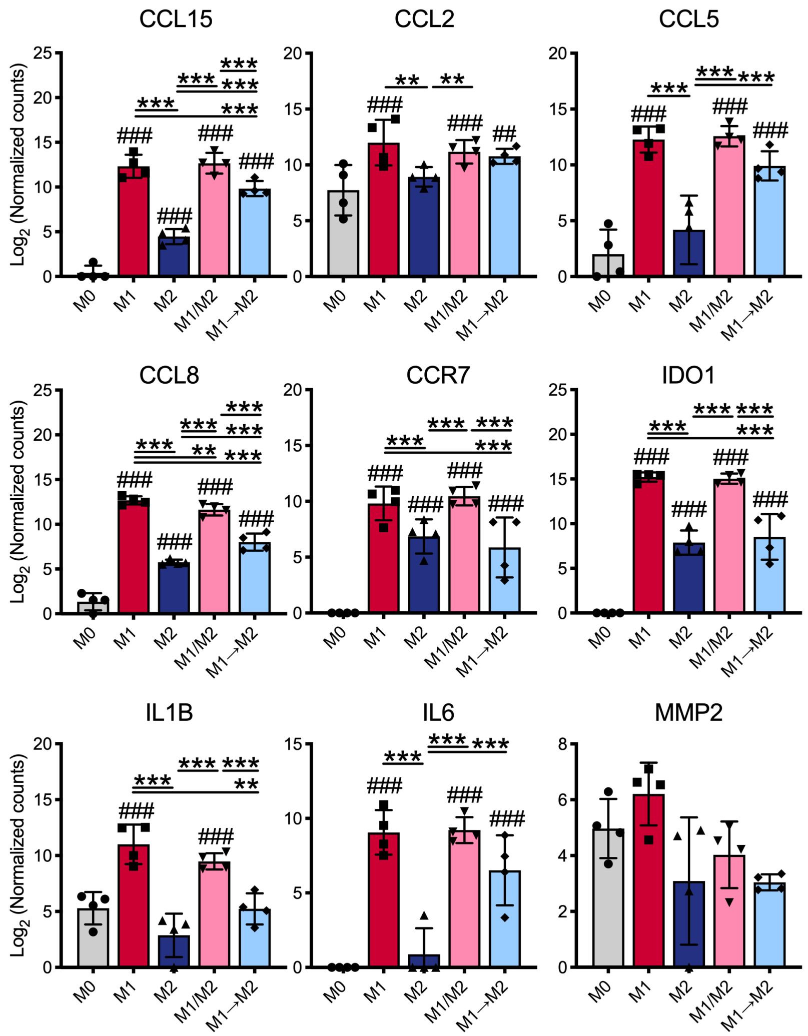


**Supplementary Figure 1**: Gene expression of M1 Markers after 72 hours of polarization analyized via NanoString. Data are represented as mean ± standard deviation. All statistical analysis was performed in GraphPad Prism 8.3 using a mixed effects analysis followed by a multiple comparisons test controlling the false discovery rate (desired rate of 0.01) using a two-stage linear step up correction method of Benjamini, Krieger, and Yekutieli **, ## p<0.01, ***, ### p<0.001, where # symbols indicate statistical significance of that data compared to M0 control.

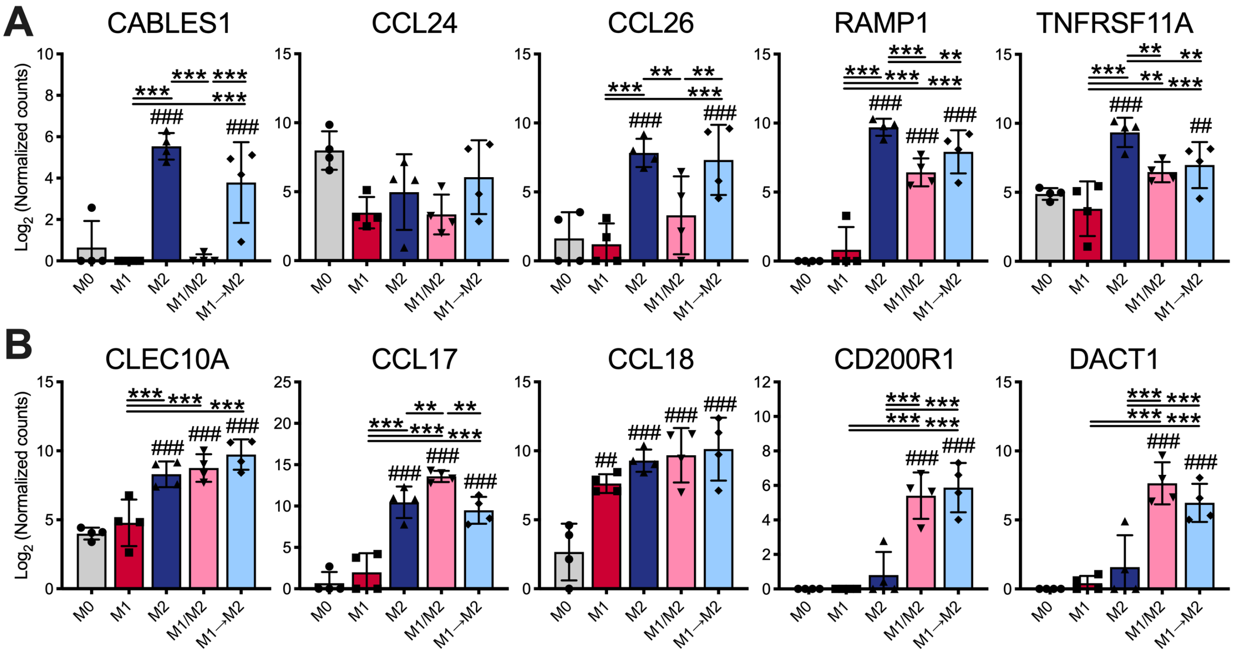


**Supplementary Figure 2**: A) Gene expression of M2 Markers after 72 hours of polarization analyized via NanoString. B) Gene expression of M2 Markers, also associated with adaptive immune function, after 72 hours of polarization analyized via NanoString. Data are represented as mean ± standard deviation. All statistical analysis was performed in GraphPad Prism 8.3 using a mixed effects analysis followed by a multiple comparisons test controlling the false discovery rate (desired rate of 0.01) using a two-stage linear step up correction method of Benjamini, Krieger, and Yekutieli **, ## p<0.01, ***, ### p<0.001, where # symbols indicate statistical significance of that data compared to M0 control.

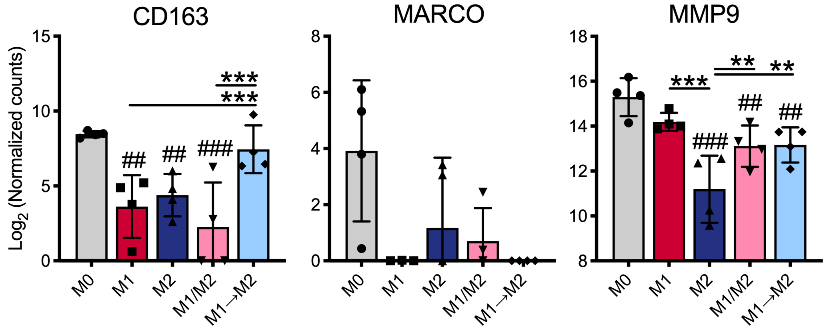


**Supplementary Figure 3**: Gene expression of M2c Markers after 72 hours of polarization analyized via NanoString. Data are represented as mean ± standard deviation. All statistical analysis was performed in GraphPad Prism 8.3 using a mixed effects analysis followed by a multiple comparisons test controlling the false discovery rate (desired rate of 0.01) using a two-stage linear step up correction method of Benjamini, Krieger, and Yekutieli **, ## p<0.01, ***, ### p<0.001, where # symbols indicate statistical significance of that data compared to M0 control.

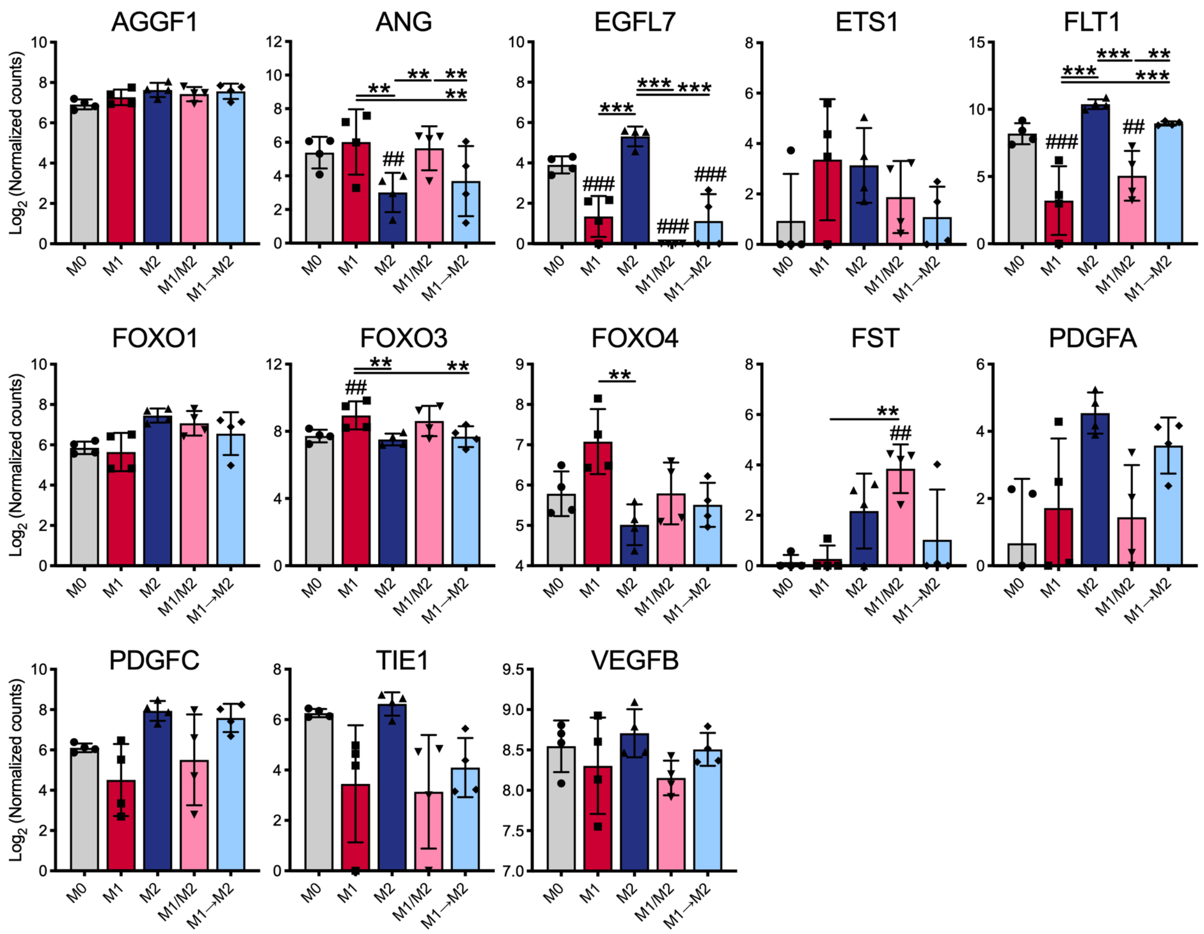


**Supplementary Figure 4**: Gene expression of Angiogenesis Markers after 72 hours of polarization analyized via NanoString. Data are represented as mean ± standard deviation. All statistical analysis was performed in GraphPad Prism 8.3 using a mixed effects analysis followed by a multiple comparisons test controlling the false discovery rate (desired rate of 0.01) using a two-stage linear step up correction method of Benjamini, Krieger, and Yekutieli **, ## p<0.01, ***, ### p<0.001, where # symbols indicate statistical significance of that data compared to M0 control.

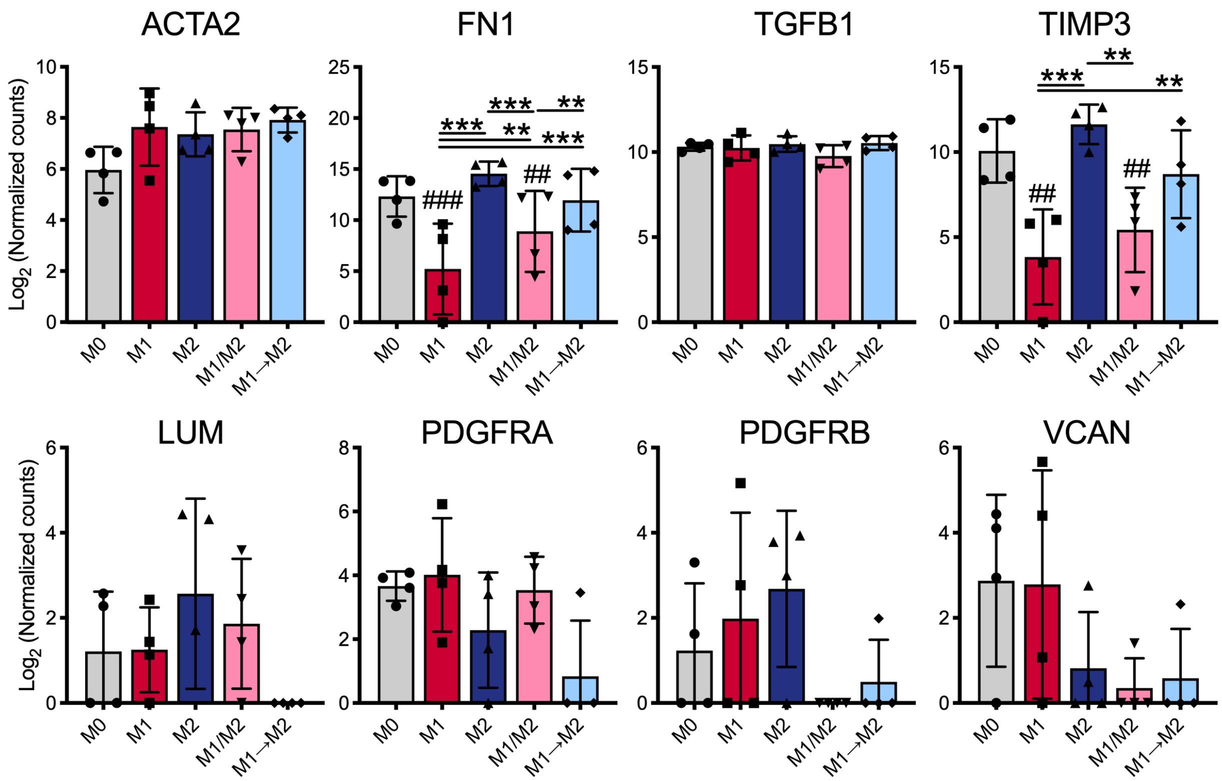


**Supplementary Figure 5:** Gene expression of ECM Markers after 72 hours of polarization analyized via NanoString. Data are represented as mean ± standard deviation. All statistical analysis was performed in GraphPad Prism 8.3 using a mixed effects analysis followed by a multiple comparisons test controlling the false discovery rate (desired rate of 0.01) using a two-stage linear step up correction method of Benjamini, Krieger, and Yekutieli **, ## $p<0.01$, ***, ### $p<0.001$, where # symbols indicate statistical significance of that data compared to M0 control.

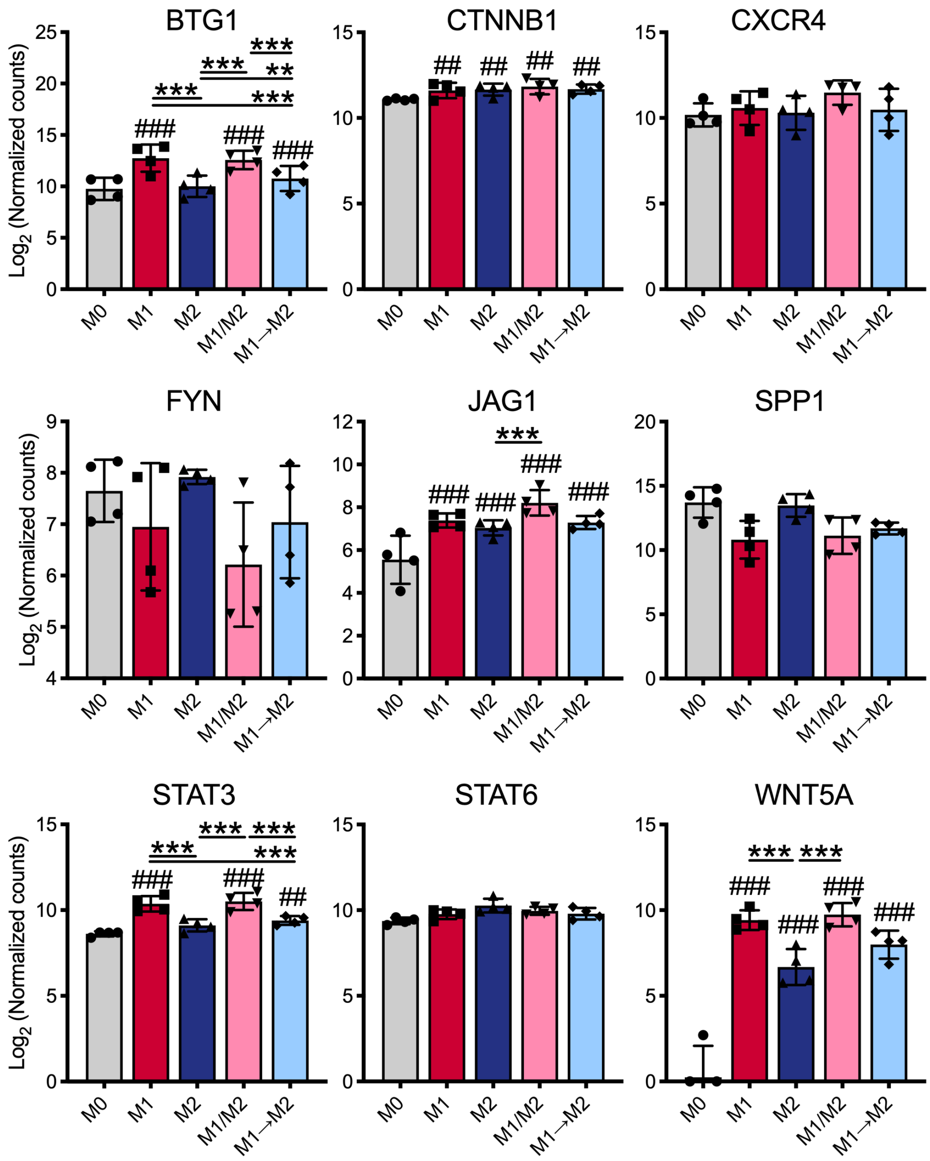


**Supplementary Figure 6**: Gene expression of Immune Signalling Markers after 72 hours of polarization analyzed via NanoString. Data are represented as mean ± standard deviation. All statistical analysis was performed in GraphPad Prism 8.3 using a mixed effects analysis followed by a multiple comparisons test controlling the false discovery rate (desired rate of 0.01) using a two-stage linear step up correction method of Benjamini, Krieger, and Yekutieli **, ## p<0.01, ***, ### p<0.001, where # symbols indicate statistical significance of that data compared to M0 control.

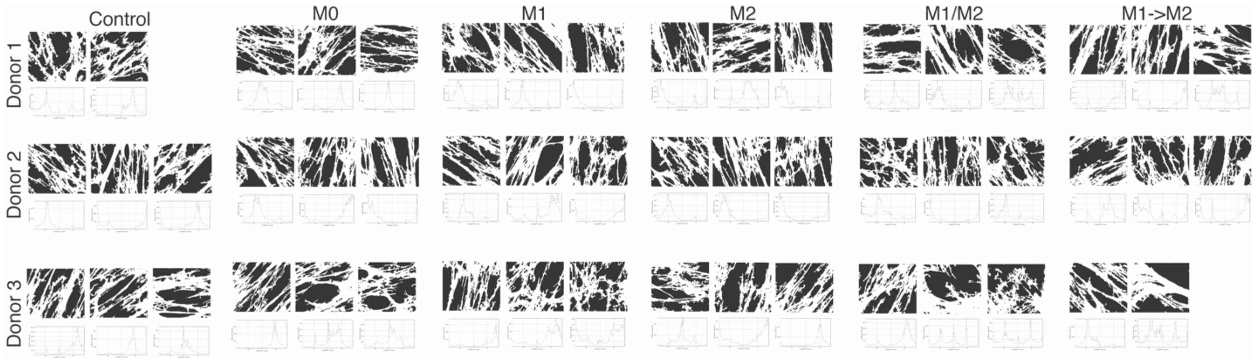


**Supplementary Figure 7**: Individual fibroblast-derived matrix segmented confocal microscopy images and corresponding Diameter J orientation plots of media-only control and primary human monocyte-derived macrophage conditioned media cultured with human dermal fibroblasts after 14 days.

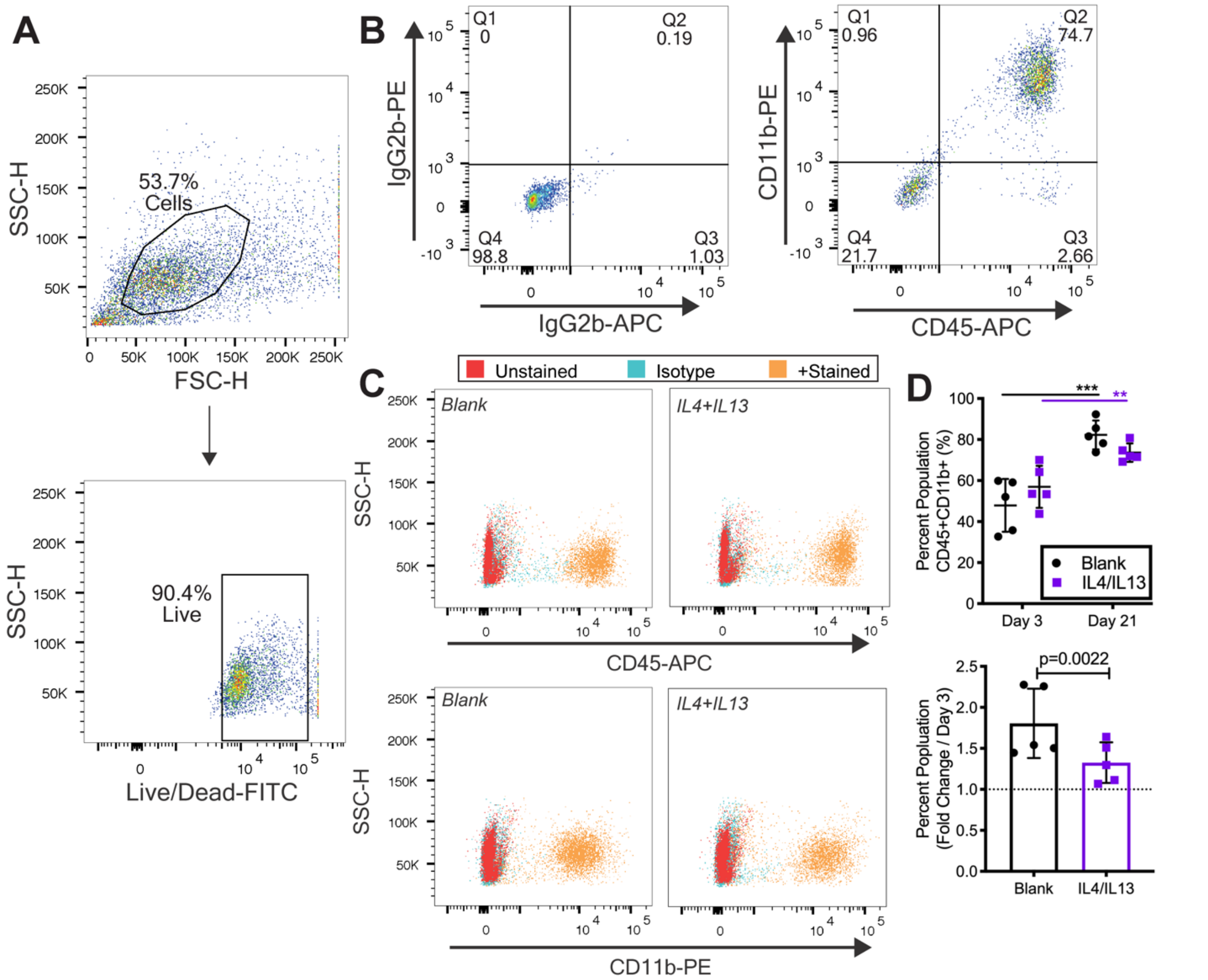


**Supplementary Figure 8**. *IL4+IL13 PLGA-GelMA hydrogels promoted a significant decrease in CD45+CD11b+ cells after 21 days in vivo*. A) Flow cytometry gating strategy for cells, live cells, and against B) isotype controls**.** C) Representative images of Blank vs. IL4+IL13 hydrogels, including unstained control, isotype control, and +stained cells for CD45 and CD11b. D) Percent population of double stained CD45+CD11b+ relative to live cells and data normalized against Day 3 timepoint. Data is represented as mean ± standard deviation. Statistical analysis was performed with a repeated measures two-way ANOVA followed by a multiple comparisons test controlling the false discovery rate using a two-stage linear step up correction method of Benjamini, Krieger, and Yekutieli or a paired two-tailed t-test $^{**}p<0.01$, $^{***}p<0.001$.

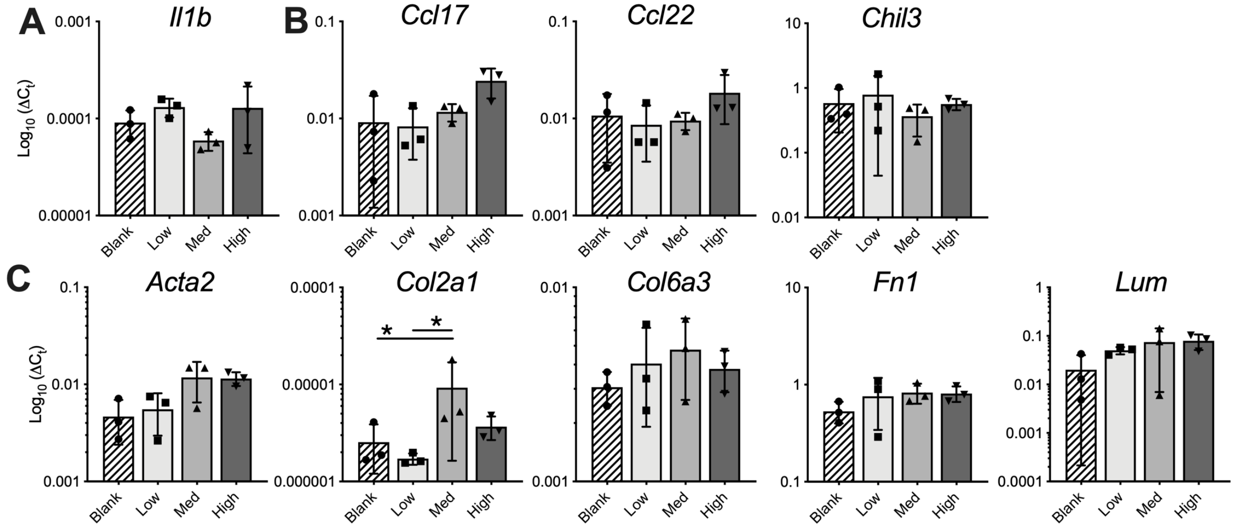


**Supplementary Figure 9**. *IL4+IL13-loaded PLGA-GelMA dose dependent effects on gene expression after 21 days in vivo*. Additional genes from pilot subcutaneous study that were not significantly different relevant to the Blank control, A) M1 Marker, B) M2 Markers, and C) ECM Markers. Data is represented as mean ± standard deviation of data normalized to housekeeping gene, *b-actin* (n=3 experimental replicates). Statistical analysis was performed using an ordinary one-way ANOVA with a Tukey's post-hoc multiple comparisons test, $^{*}p<0.05$.

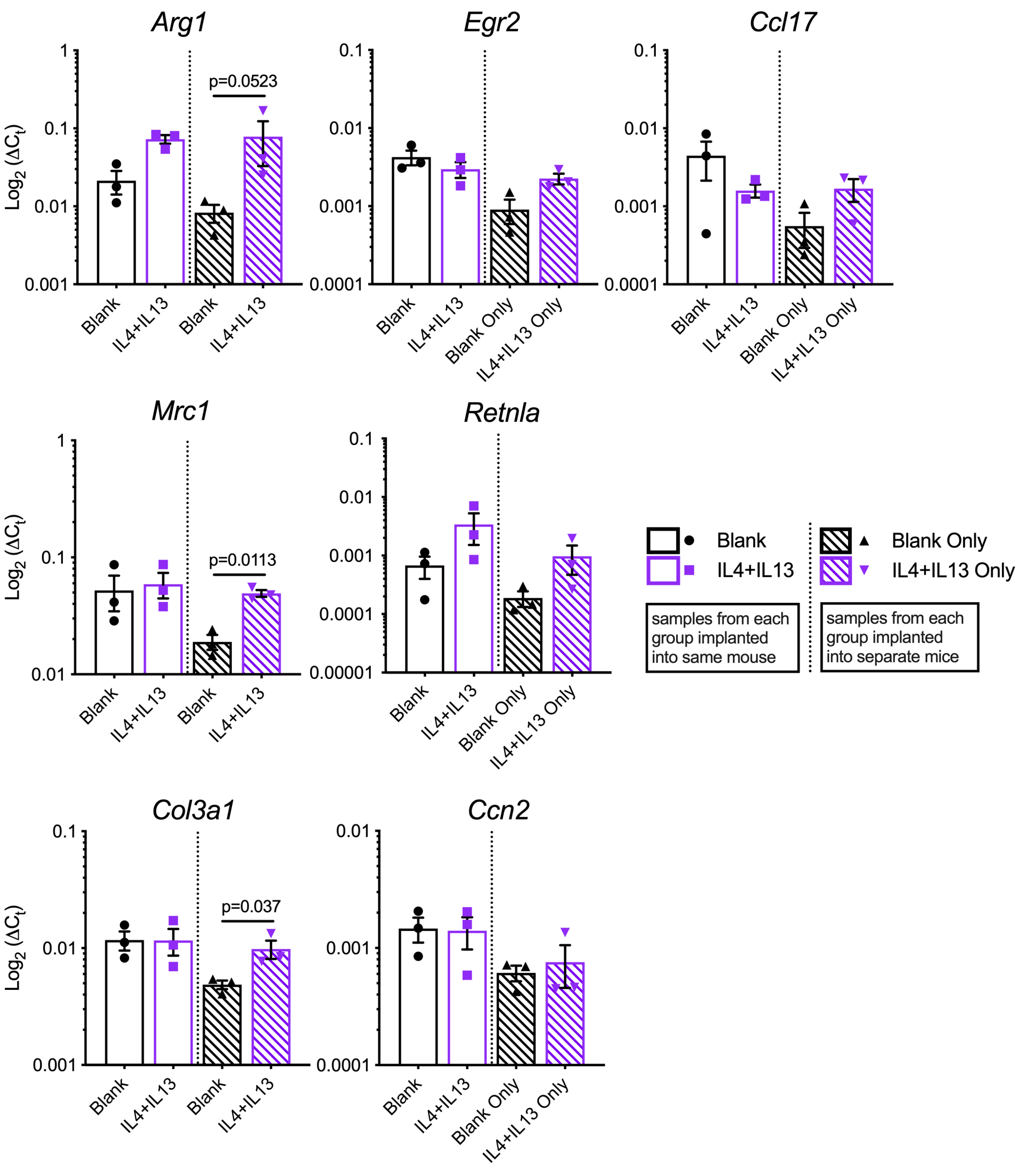


**Supplementary Figure 10**. *Hydrogels implanted into separate mice, rather than within the same mouse, promote differences in gene expression of heterogenous tissue explants after 3 days in vivo*. Data represented as mean ± standard error of the mean. Data was normalized to housekeeping gene, *b-actin*. Statistical analysis for the samples within same mouse was performed with a paired two-tailed t-test, while analysis for samples within individual mice was performed using unpaired two-tailed student's t-test with a Welch's correction.

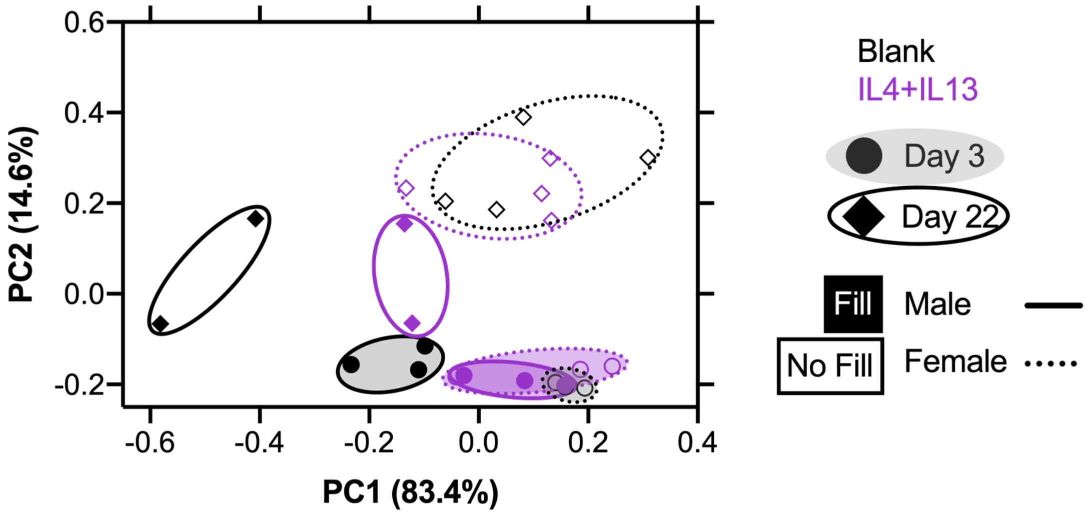


**Supplementary Figure 11**: Principal component analysis reveals that 99% of the variance derives from time and group (Blank vs. IL4+IL13), but also separated by sex.

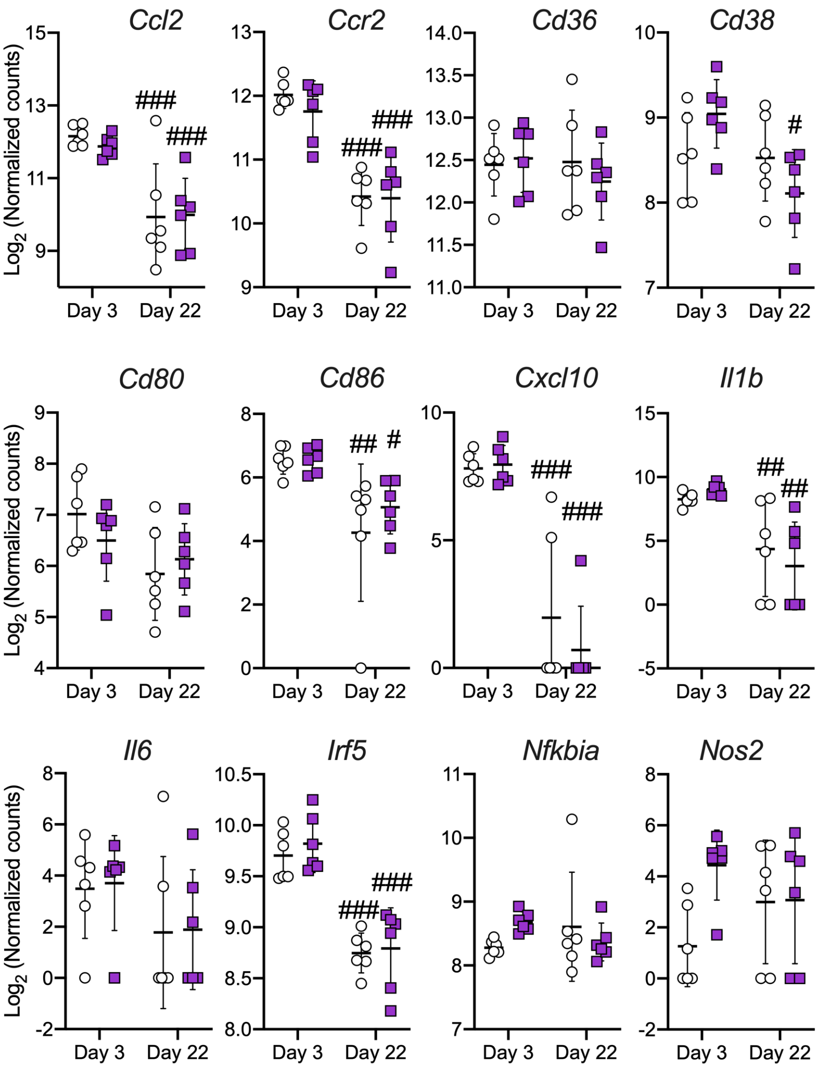


**Supplementary Figure 12**: Individual gene expression analysis of M1 markers performed via NanoString of IL4+IL13 PLGA-GelMA hydrogels explanted from subcutaneous space after 3 and 22 days *in vivo*. # p<0.05, ## p<0.01, ### p<0.001, where # symbols indicate statistical significance of that data compared to corresponding Day 3 data. Data represented as mean ± standard deviation, n=6 mice per group, per time point.

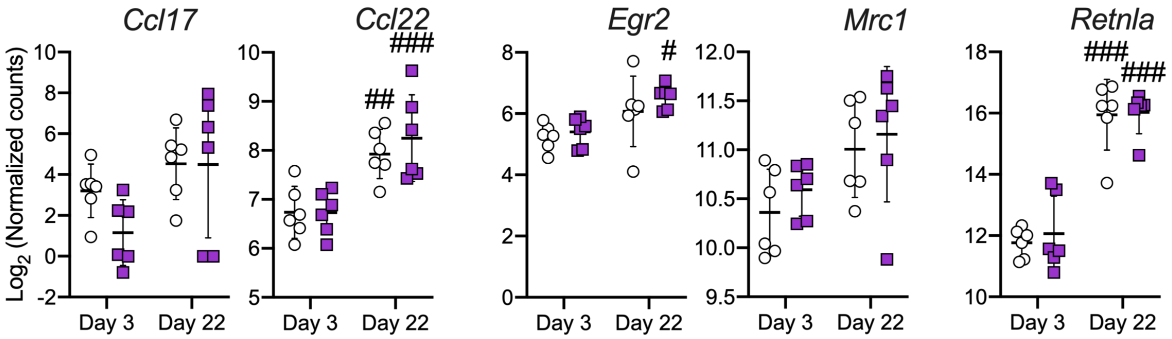

**Supplementary Figure 13**: Individual gene expression analysis of M2 markers performed via NanoString of IL4+IL13 PLGA-GelMA hydrogels explanted from subcutaneous space after 3 and 22 days *in vivo*. # p<0.05, ## p<0.01, ### p<0.001, where # symbols indicate statistical significance of that data compared to corresponding Day 3 data. Data represented as mean ± standard deviation, n=6 mice per group, per time point.

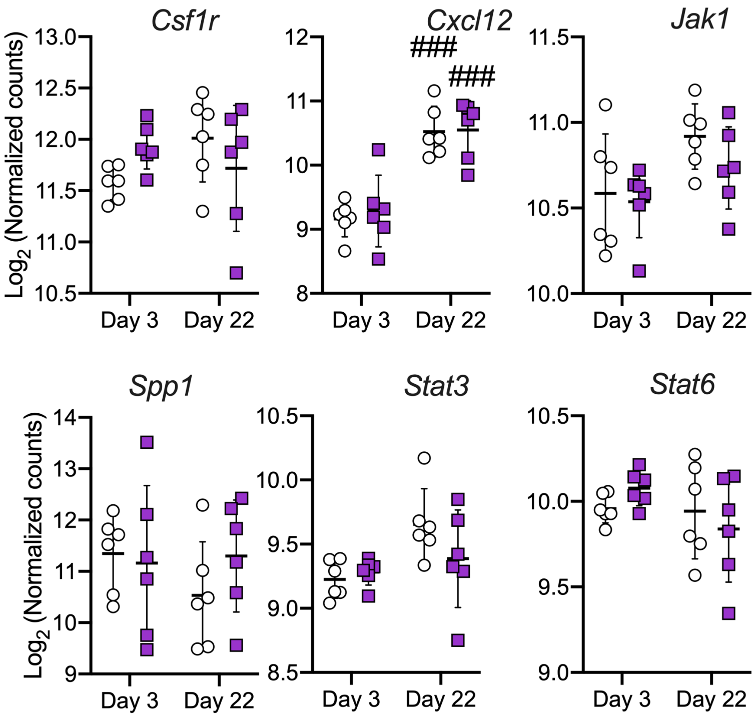


**Supplementary Figure 14**: Individual gene expression analysis of Immune Signaling markers performed via NanoString of IL4+IL13 PLGA-GelMA hydrogels explanted from subcutaneous space after 3 and 22 days *in vivo*. ### p<0.001, where # symbols indicate statistical significance of that data compared to corresponding Day 3 data. Data represented as mean ± standard deviation, n=6 mice per group, per time point.

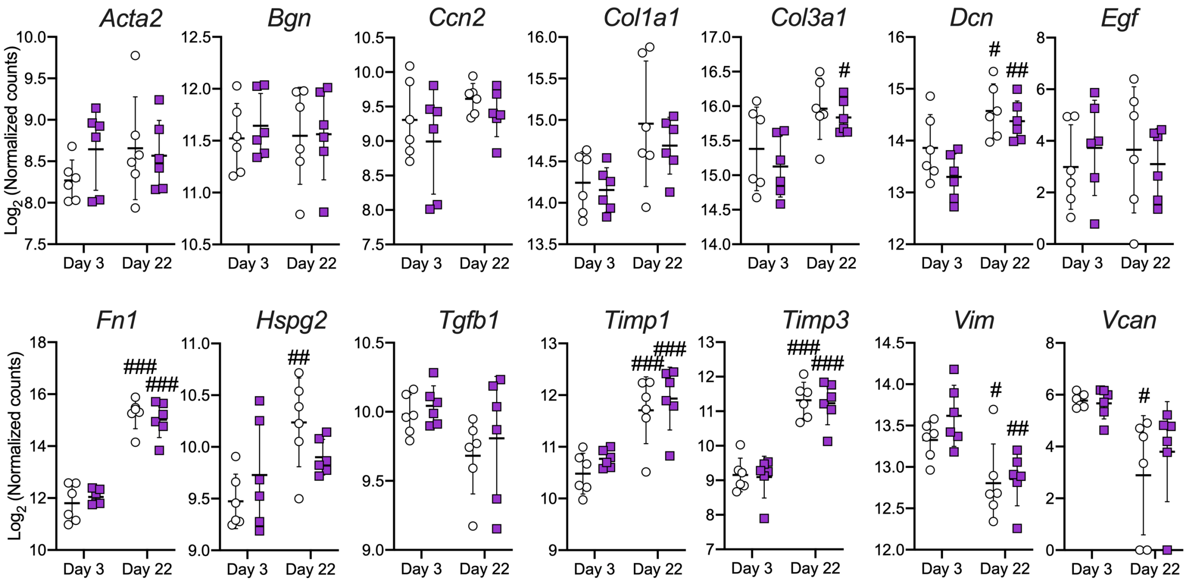


**Supplementary Figure 15**: Individual gene expression analysis of ECM markers performed via NanoString of IL4+IL13 PLGA-GelMA hydrogels explanted from subcutaneous space after 3 and 22 days *in vivo*. # p<0.05, ## p<0.01, ### p<0.001, where # symbols indicate statistical significance of that data compared to corresponding Day 3 data. Data represented as mean ± standard deviation, n=6 mice per group, per time point.

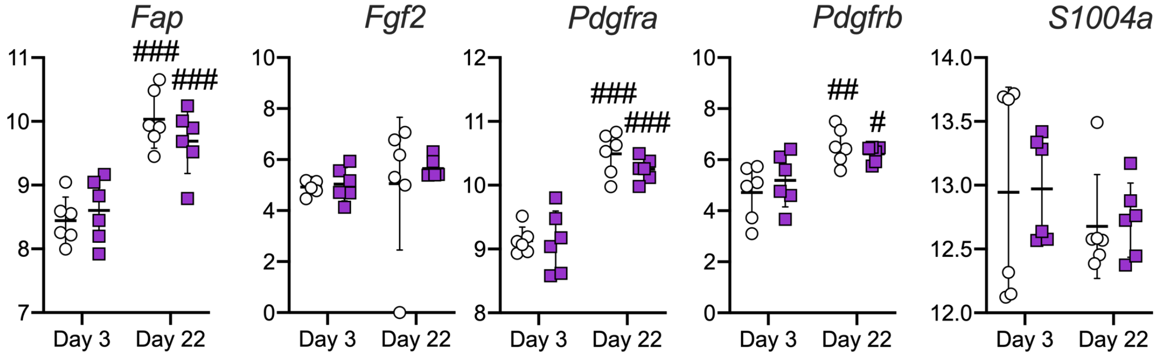


**Supplementary Figure 16**: Individual gene expression analysis of Fibroblast markers performed via NanoString of IL4+IL13 PLGA-GelMA hydrogels explanted from subcutaneous space after 3 and 22 days *in vivo*. # p<0.05, ## p<0.01, ### p<0.001, where # symbols indicate statistical significance of that data compared to corresponding Day 3 data. Data represented as mean ± standard deviation, n=6 mice per group, per time point.

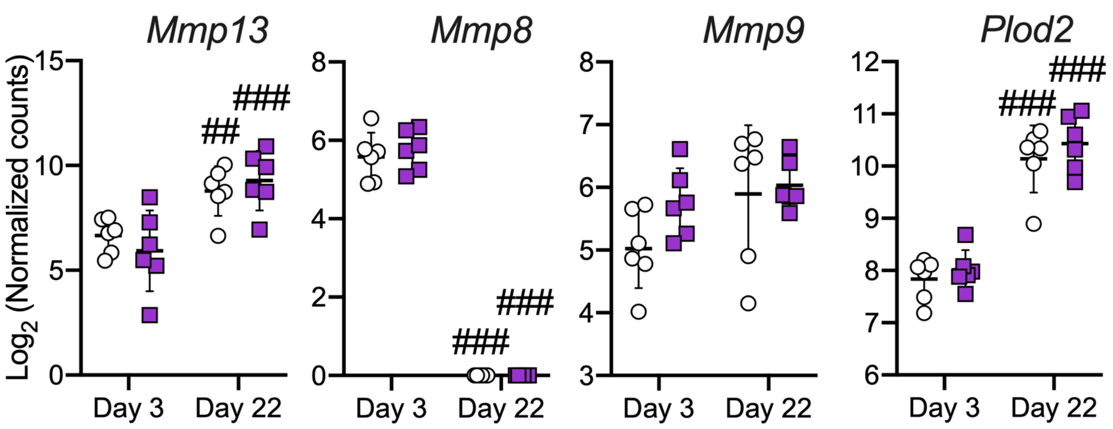


**Supplementary Figure 17**: Individual gene expression analysis of Protease markers performed via NanoString of IL4+IL13 PLGA-GelMA hydrogels explanted from subcutaneous space after 3 and 22 days *in vivo*. # $p<0.05$, ## $p<0.01$, ### $p<0.001$, where # symbols indicate statistical significance of that data compared to corresponding Day 3 data. Data represented as mean ± standard deviation, n=6 mice per group, per time point.

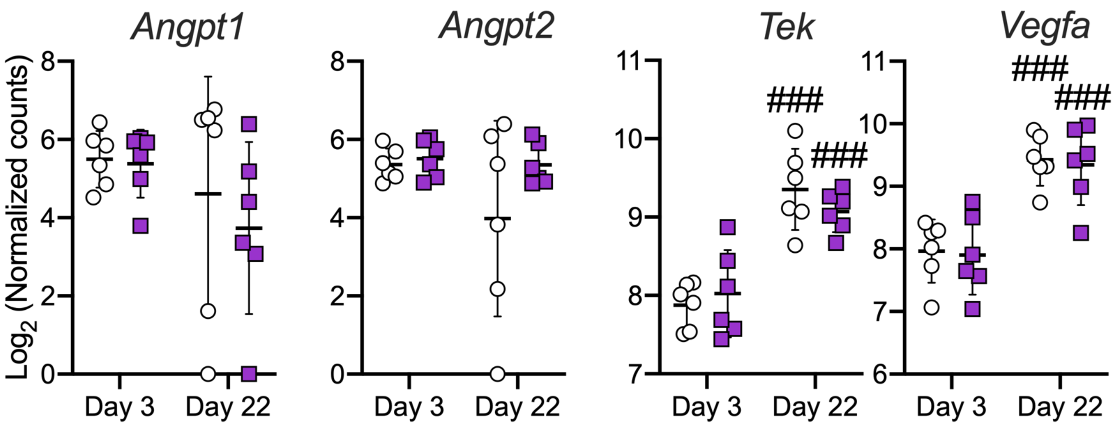


**Supplementary Figure 18**: Individual gene expression analysis of Angiogenesis markers performed via NanoString of IL4+IL13 PLGA-GelMA hydrogels explanted from subcutaneous space after 3 and 22 days *in vivo*. # $p<0.05$, ## $p<0.01$, ### $p<0.001$, where # symbols indicate statistical significance of that data compared to corresponding Day 3 data. Data represented as mean ± standard deviation, n=6 mice per group, per time point.

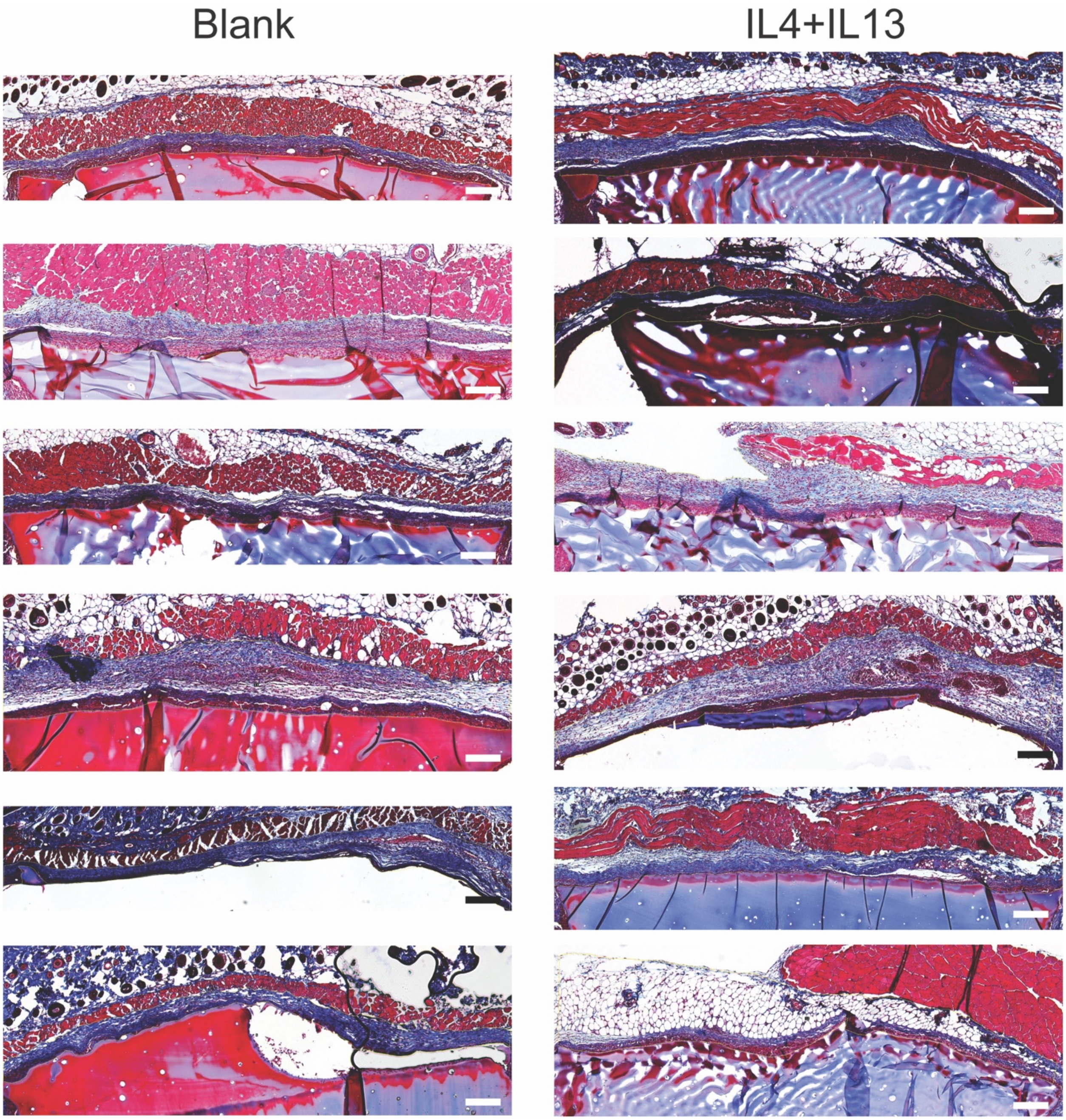


**Supplementary Figure 19**: All hydrogel explants (attached to skin) stained Masson's Trichrome, imaged, and fibrous capsule manually traced with a thin yellow line in ImageJ. Scale bar = 100μm.

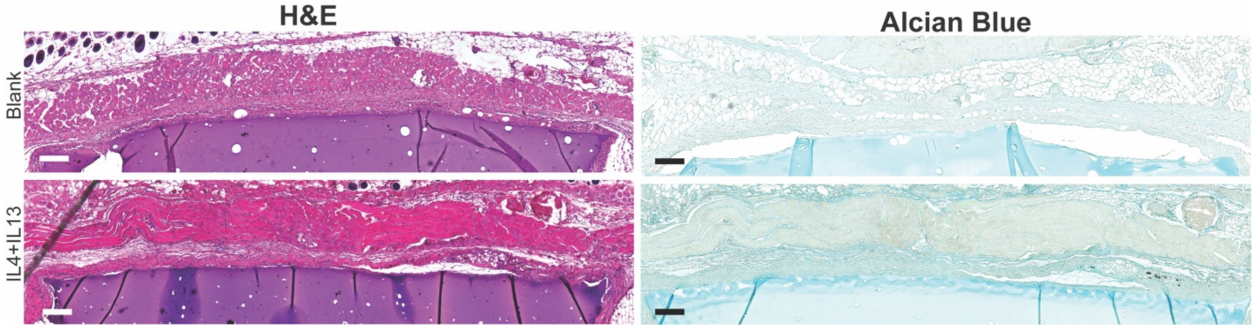


**Supplementary Figure 20**: Representative images of hemotoxylin and eosin (H&E) and Alcian Blue histological staining. Scale bar = 100μm.

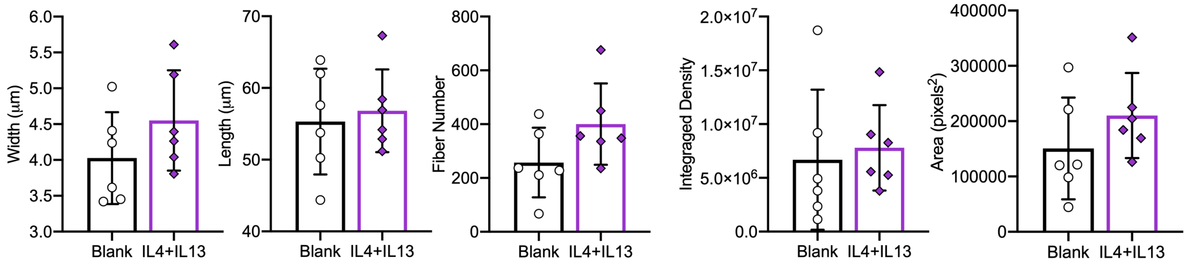


**Supplementary Figure 21**: Additional SHG analysis metrics. Data represented as mean ± standard deviation, n=6 mice per group.
